# Identification and characterization of four bacteriome- and mycobiome-derived subtypes in tumour and adjacent mucosa tissue of colorectal cancer patients

**DOI:** 10.1101/2022.09.27.509391

**Authors:** Manuela Salvucci, Liam Poynter, Reza Minerzami, Steven Carberry, Robert O’Byrne, James Alexander, Diether Lambrechts, Kirill Veselkov, James Kinross, Jochen H. M. Prehn

**Affiliations:** Centre for Systems Medicine, Department of Physiology and Medical Physics, Royal College of Surgeons in Ireland, Dublin, Ireland; Department of Surgery and Cancer, Imperial College London, London, United Kingdom; Department of Colorectal Surgery, The Royal Free Hospital, Hampstead, London, United Kingdom; Department of Metabolism, Digestion and Reproduction, Imperial College London, London, United Kingdom; VIB Center for Cancer Biology, VIB, Leuven, Belgium

**Keywords:** colorectal cancer, microbiome, bacteriome, bacteria, mycobiome, fungi, subtyping, signatures, prognostic biomarker

## Abstract

**Objective:** Here, we systematically investigated alterations in the bacteriome and mycobiome of CRC patients in tumours and matched adjacent mucosa resulting in the identification of microbiome-based subtypes associated with host clinico-pathological and molecular characteristics.

**Design:** Diversity and composition of bacteriome and mycobiome of tumour and adjacent mucosa, resulting subtypes were computationally deconvoluted from RNA sequencing, using >10000 samples from in-house and publically available patient cohorts.

**Results:** The bacteriome of tumours had higher dominance and lower α-diversity compared to matched adjacent local and distant mucosa. Tumours were enriched with *Proteobacteria* (*Gammaproteobacteria* class), *Fusobacteria* (including *Fusobacterium Nucleatum* species) and *Basidiomycota* fungi (*Malasseziaceae* family). Tumours were depleted of *Bacteroidetes* (*Bacteroidia* class), *Firmicutes* (*Clostridia* class) and *Ascomycota* (*Sordariomycetes* and *Saccharomycotina*). Tumours and adjacent mucosa samples were classified into 4 microbial subtypes, termed C1 to C4, based on the bacteriome and mycobiome composition. The bacterial *Propionibacteriaceae*, *Enterobacteriaceae*, *Fusobacteriaceae*, *Bacteroidaceae* and *Ruminococcaceae* and the fungal *Malasseziaceae*, *Saccharomycetaceae* and *Aspergillaceae* were among the key families driving the microbial subtyping. Microbial subtypes were associated with distinct tumour histology and patient phenotypes and served as an independent prognostic marker for disease-free survival. Key associations between microbial subtypes and alterations in host immune response and signalling pathways were validated in the TCGA pan-cancer cohort. The microbial subtyping demonstrated stratification value in the pan-cancer settings beyond merely representing differences in survival by cancer type.

**Conclusions:** This study demonstrates the translational potential of microbial subtyping in CRC patient stratification, and provides avenues to design tailored microbiota modulation therapy to further precision oncology.

**Statement of significance:** *What is already known on this subject?:*

- The microbiome has been implicated in the pathogenesis, progression and therapeutic response in patients diagnosed with CRC and other cancers.
- The vast majority of studies to date has focussed on investigating the bacteriome while the critical role played by the mycobiome in shaping cancer has begun to be explored more recently.
- The bacterial and fungal composition and diversity in on-tumour tissue compared to matched local and distal off-tumour mucosa is largely unexplored.
- Tumorigenesis may be promoted via alterations of the microbiome ecosystem that may be better recapitulated by a multi-kingdom microbial signature rather than by the abundance of individual bacterial or fungal microorganisms.

*What are the new findings?:*

- On-tumour tissue was enriched with *Fusobacteria, Proteobacteria, Basidyomicota* and depleted of *Bacteroidetes, Firmicutes, Ascomycota* compared with adjacent off-tumour mucosa.
- We stratified >600 CRC patients into four distinct microbial-based subtypes (C1-C4) according to their bacteriome and mycobiome composition. The microbial subtypes were associated with distinct prognosis, clinical phenotypes such as staging, tumour location, history, lymphovascular invasion, TP53 status and clinical outcome.
- Furthermore, the majority of matched adjacent mucosa samples were classified as C1 and paired tumour-matched normal samples demonstrated a robust shift towards the C1 subtype in off-tumour tissue, suggesting that the C1 subtype may recapitulate a healthier-like microbiome. This hypothesis was supported by the microbial subtyping of colon samples from healthy subjects that were categorised as C1 almost exclusively.
- The identified microbial subtyping demonstrated stratification value in the pan-cancer settings (n=28 additional solid cancer indications spanning >9000 samples) beyond merely representing differences in survival by cancer type, providing the strongest stratification in liver cancer.

*How might it impact on clinical practise in the foreseeable future?:*

- This study laids the foundation to develop a microbial signature as biomarker to clinically manage CRC and other solid cancers and potentially lead to microbiome-based companion diagnostics to design microbiota modulation therapy tailored to specific patients subgroups with distinct bacteriomes and mycobiomes.

## Introduction

The bacteriome and mycobiome, collectively referred to as the microbiome, is a key player in CRC pathogenesis, progression and response to chemotherapy, radiotherapy and immunotherapy [1–9]. Several research groups, including the present authors’, have focused primarily on the bacteriome, leading to the identification of specific species involved in carcinogenesis such as *Fusobacterium, Escherichia Coli* and *Bacteroides fragilis* [9–13]. More recently, alterations in fungal taxa, such as *Malassezia, Saccharomycetes* and a shift in *Ascomycota:Basidiomycota* ratio, have been implicated as cancer-disease modifiers [14–16].

Microbial modulation and exacerbation in CRC is likely attributable to alterations of composition and diversity of the microbiome rather than to a single microbial species [17]. These alterations in microbial signatures may include a decrease in diversity induced by a decrease in beneficial species coupled with an outgrowth of pathogens. Microbial species may drive CRC by interacting with host cell via invasion and translocation along with the creation of biofilms, by modulating the host immune response and signalling pathway, activating EMT transition and cell migration, and by secreting metabolites and toxigenic molecules [1,3,4,7,11,18]. Microbiome characterizations to date in CRC have largely focused on tumour tissue alone. Recent research has highlighted how the mucosal tissue surrounding tumours may be substantially different from “truly’’ healthy tissue excised from non-tumour bearing subjects [19]. Slaughter and collaborators coined the term “field cancerization” describing the adjacent mucosa as an intermediate state featuring epithelial cells and tumour microenvironment (TME) with normal morphological characteristics, but altered signalling [20].

Thus, in this study, we performed a large-scale unbiased characterization of the bacteriome and mycobiome in tumour and matched tumour-adjacent and tumour-remote mucosa, using an in-house CRC cohort as well as external, deeply-annotated and clinically relevant colorectal cohorts. We investigated, for the first time to our knowledge, the association between microbial-based subtypes with clinical and phenotypical manifestations of CRC. The impact of composition and diversity of the host microbiome is not limited to CRC [21], but has been implicated in progression and therapeutic resistance in other cancers [22–26]. Thus, we additionally extended this analysis to 28 other solid cancer types to evaluate the applicability of the microbial subtyping beyond the CRC settings.

## Results

We characterised the composition of the bacteriome and mycobiome in tumours and matched normal mucosa of CRC patients and determined the association of these profiles with clinical and phenotypic CRC manifestations. We estimated bacterial and fungal composition from RNA sequencing experiments using a validated subtractive method (*PathSeq* [27]) in in-house and publicly available cohorts, (**Sup. Table 1**). We evaluated the quality of the microbial estimates with two sets of orthogonal analysis determining i) the agreement with composition from an independent method and ii) retrieval of expected clinically-relevant microbial signatures.

### Bacterial composition deconvoluted from transcriptomics recapitulates expected clinically-relevant microbial profiles

We performed RNA sequencing on an in-house cohort of tissue samples obtained from n=26 CRC patients accrued at St. Mary’s Hospital (Imperial College London, London, UK), referred to as “ICL” cohort. The cohort was largely composed of male (70%) patients diagnosed with stage II-III (78%) cancer of the colon (78%) with 43% of the cases presenting with lymphovascular invasion. Clinical, demographic and pathological characteristics of the patients included in the ICL cohort is reported in **Sup. Table 2**. We collected fresh-frozen samples from the tumour bulk and matched normal mucosa at 5 cm and 10 cm from the tumour margins to additionally investigate the microbiome composition in local and distant spatial locations. We had previously determined [1] bacterial composition in a subset of the ICL cohort patients (n=18/26, 69%) by 16S rRNA. We compared the bacterial phyla composition determined by 16S rRNA against the estimates deconvoluted from RNA sequencing pipeline used in this study (*PathSeq*) (“ICL cohort”, n=49 samples from n=26 patients) on the samples assayed by both methods. The 16S rRNA identified n=19 unique phyla across all samples, 18 of which, (in yellow in **Fig. 1A**), were also identified by the *PathSeq* pipeline. The *PathSeq* pipeline identified additional low abundance taxa compared to the 16S rRNA method, totalling n=36 unique phyla. When comparing phylum abundance, we observed good agreement, both overall and within samples collected from bulk tumour and matched tissue resected at 5 cm and 10 cm from the the tumour margins (P<0.0001; **Fig. 1B**).

**Figure 1.**
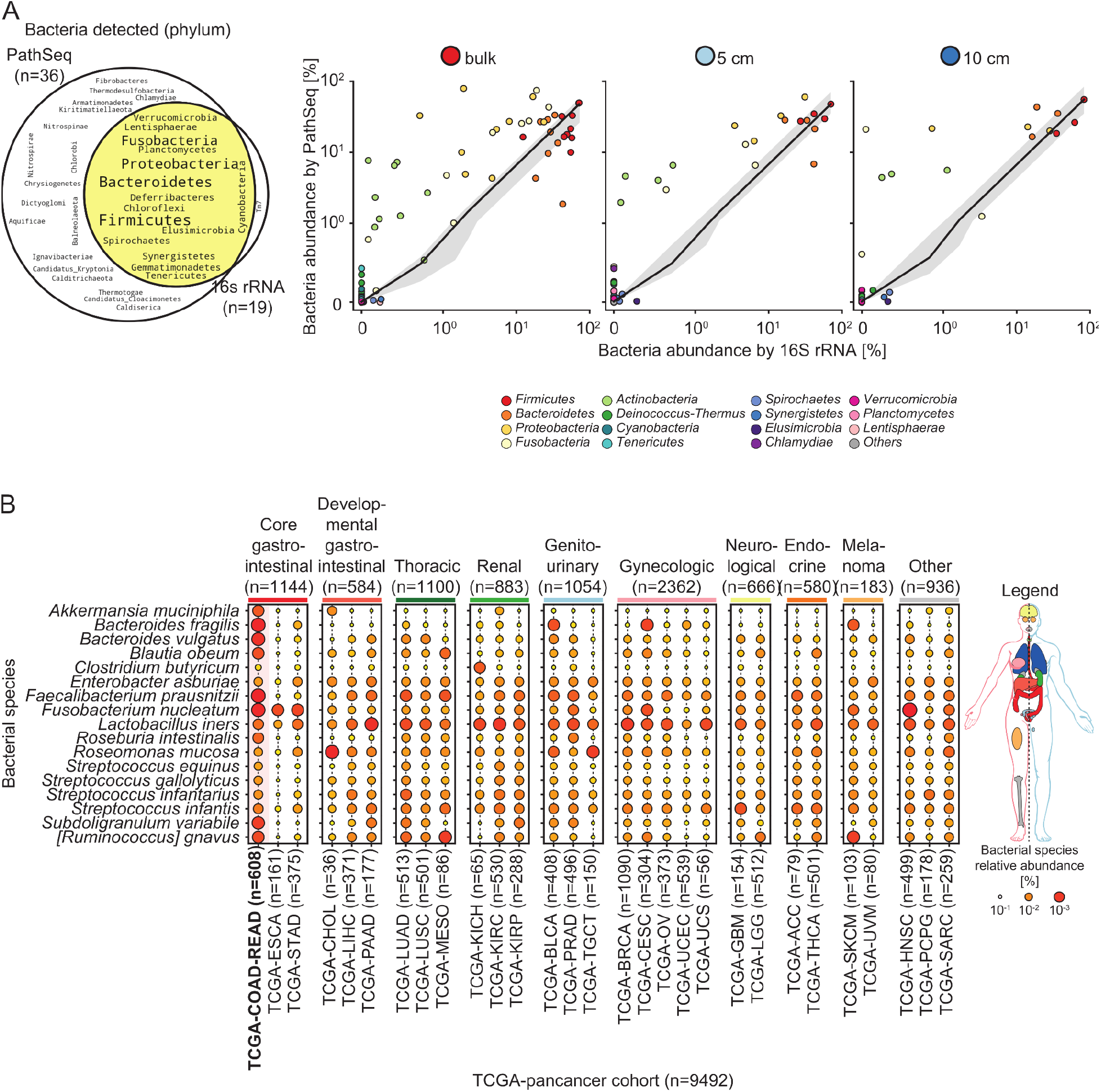
Bacterial composition deconvoluted from transcriptomics tally with estimates from an independent gold-standard method in the in-house ICL cohort (A) and recapitulates expected clinically-relevant microbial profiles in the TCGA pan-cancer cohort (B). **A.** Benchmark of the bacterial composition at the phylum taxonomic rank determined using the gold standard 16S rRNA against the estimates deconvoluted from RNA sequencing experiments with the PathSeq pipeline on tissue samples resected from the bulk tumour and local (5 cm) and distant (10 cm) adjacent mucosa in the in-house ICL cohort (n= 49 samples from n=26 patients). **Left hand-side.** Venn diagram indicates the number and name of bacterial phyla detected by either or both (highlighted in yellow) workflows in at least a sample. The average phyla relative abundance is encoded by the font size with larger fonts indicating higher bacterial presence. **Right hand-side.** Scatter plot detailing the relationship between relative abundance detected by 16s rRNA with PathSeq-derived estimates by bacterial phylum and tissue sampling site (bulk on-tumour vs. off-tumour adjacent mucosa at 5 and 10 cm). **B.** Benchmark of the microbial composition determined using the PathSeq pipeline against expected and clinically-relevant bacterial species profiles in tumour samples from patients diagnosed with CRC (TCGA-COAD-READ, n=608, highlighted in red) and other solid oncological indications (n=28 additional cancer types, totalling n=9492 patients). Selected bacterial species robustly reported in the literature as either being constituents of the native colorectal commensal community or putative cancer-drivers or disease-modifiers.Average species relative abundance by cancer type is colour- and size-encoded with darh=ker and bigger markers indicating higher bacterial species presence. Clinically-related cancers were grouped and colour-coded as indicated in the legend.

As a further quality control, we benchmarked the microbial composition determined using the *PathSeq* pipeline against expected and clinically-relevant microbial profiles reported in the literature. We determined the microbial composition from RNA sequencing experiments of patients of The Cancer Genome Atlas (TCGA) collection including both samples from patients diagnosed with colorectal cancer (TCGA-COAD-READ, n=608) along with tumour resections of subjects diagnosed with other solid oncological indications (n=28 additional cancer types, totaling n=9492 patients). Clinical, demographic and pathological characteristics of the cancer patients for the colorectal and pan-cancer cohorts are summarised in **Sup. Table 2** and **Sup. Table 3**, respectively. We selected bacterial species robustly reported in the literature as either being constituents of the native colorectal commensal community or putative cancer-drivers or disease-modifiers. We hypothesised that a robust pipeline would i) recover bacterial species known to be constituents of the gut microbiome and would ii) yield higher average relative abundance (RA) scores in CRC patients compared to other cancer types, particularly for species linked specifically to colorectal cancer. Indeed, we identified all bacterial species we set out to investigate in the TCGA samples. In line with literature evidence, we confirmed that the vast majority of the selected species were found in higher abundance in primary tumours of colorectal cancer patients (TCGA-COAD-READ) compared to those from other core and developmental gastrointestinal cancers and other oncological indications (**Fig. 1C)**. This subset included species such as *Blautia_obeum, Faecalibacterium_prausnitzii, Roseburia_intestinalis, Subdoligranulum_variabile*, and *[Ruminococcus]_gnavus* [2,21,28] found in the gastrointestinal flora and markers of dysbiosis, *Bacteroides_fragilis* and *Bacteroides_vulgatus* associated with promoting inflammation and cancer development, *Akkermansia_muciniphila* reported to modulate response to treatment chemotherapy [29] and immunotherapy [30], *Fusobacterium_nucleatum* associated with distinct molecular characteristics and patients prognosis in CRC tumours [1,10–13,31–34].

### Higher dominance and lower diversity in on-tumour samples compared to off-tumour adjacent mucosa in the bacteriome, but not the mycobiome of CRC patients

Having established the applicability of this computational pipeline, we next compared the diversity of the bacteriome and mycobiome from tumour and matched normal tissue in CRC patients of the ICL and the TCGA-COAD-READ cohorts (**Fig. 2**). We characterised the within-sample α-diversity and across-samples β-diversity by computing multiple metrics from phylum taxa. We found no statistically significant difference in the number of bacterial and fungal phyla observed in tumour samples vs. matched normal tissue in neither cohorts, s indicated by the α-diversity Chao index (**Fig. 2A.1-2-B.1-2**, **Sup. Fig. 1A.1-B.1**). We observed an increase in α-diversity dominance and reduced Shannon and Simpson E indices when comparing tumour with local (5 cm) and distant (10 cm) matched normal tissue in the ICL cohort (**Fig. 2A.1**). These findings were confirmed when comparing tumour and matched normal tissue in the ICL cohort (**Sup. Fig. 1**) and TCGA-COAD-READ cohorts (**Fig. 2A.2**). In contrast, no statistically significant differences in any of the α-diversity indices tested were observed when comparing the mycobiome of tumours with local and distant matched normal tissue in neither the in-house ICL nor the TCGA cohorts (**Fig. 2B**, **Sup. Fig. 1**).

**Figure 2.**
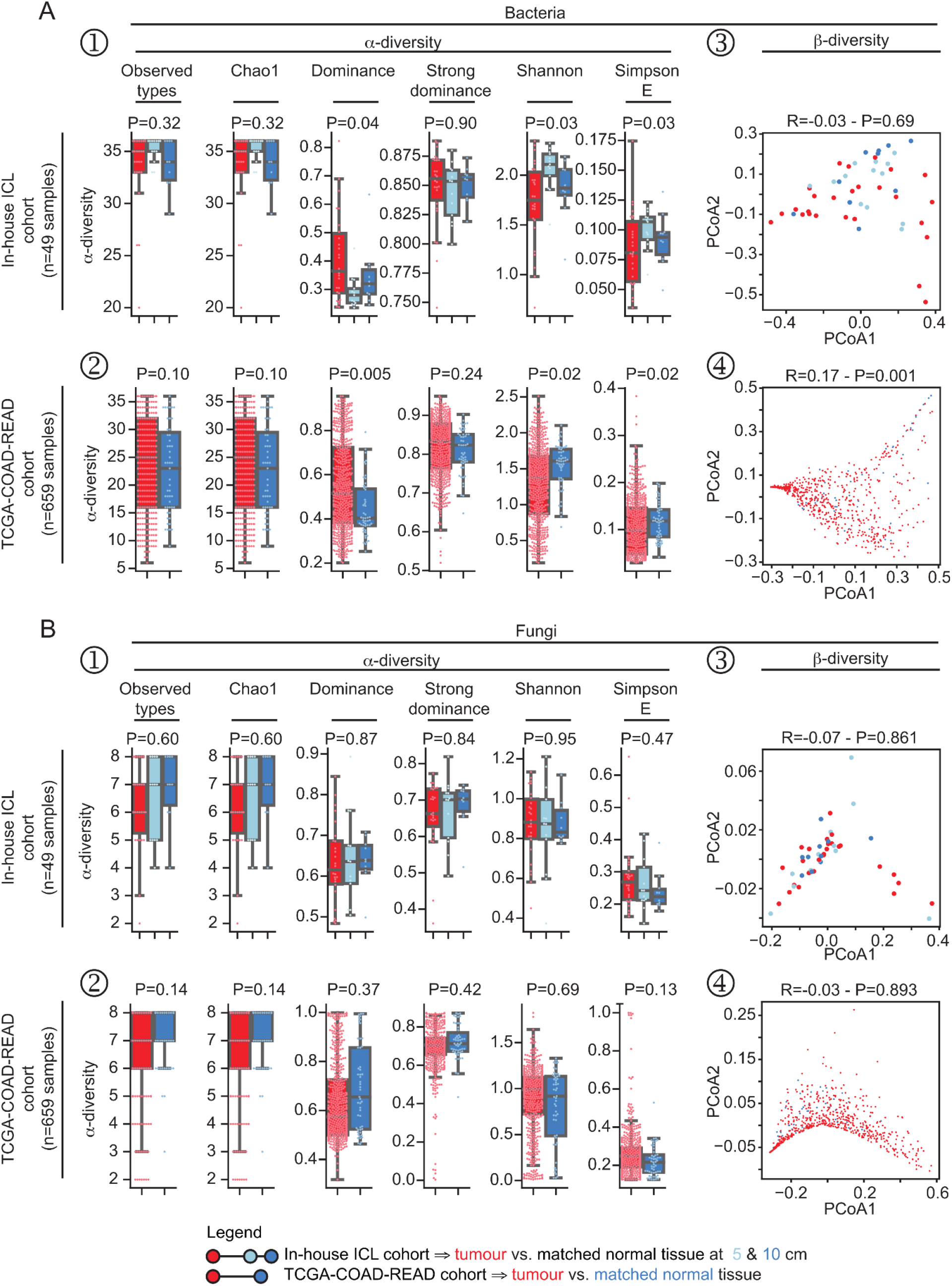
a- and β-diversity metrics for bacterial and fungal phyla in on-tumour tissue samples compared to off-tumour adjacent mucosa in tissue resections from CRC patients. **A-B.** Within-sample αdiversity (**A.1-2, B.1–2**) and across-samples β-diversity (**A.3-4, B.3-4**) metrics comparing bacterial (**A**) and fungal (**B**) ecological scores in on-tumour and off-tumour adjacent mucosa samples from patients of the in-house ICL and TCGA-COAD-READ cohorts. α-diversity indices included observed types, Chao1, (strong)-dominance, Shannon and Simpson E. β-diversity was quantified by performing unsupervised principal component analysis on Bray-Curtis distances. Statistical significance differences between β-diversity between on-tumour vs. off-tumour adjacent mucosa was performed using analysis of similarities (ANOSIM).

We investigated inter-sample diversity as measured by the Bray-Curtis β-diversity index. In line with previous literature, we identified a trend, albeit not statistically significant, whereby ICL tumours had a distinct composition from matched normal tissue, either when analysing separately (**Fig. 2A.3**) or combining (**Sup. Fig. 1A.2**) samples collected at 5/10 cm from the tumour margins. We corroborated these findings in the TCGA-COAD-READ cohort (**Fig. 2A.4**). In contrast, we observed no statistical significant difference in Bray-Curtis β-diversity of the mycobiome in either cohorts (**Fig. 2B.3-4**, **Sup. Fig. 1**).

### Distinct enrichment and depletion of bacteria and fungi between on-tumour and off-tumour adjacent mucosa

Next, we assessed the bacterial and fungal composition in tumours and matched normal tissue of the ICL and the TCGA-COAD-READ cohorts (**Fig. 3A**). In line with the literature, the most abundant phyla identified in both cohorts were *Actinobacteria, Bacteroidetes, Firmicutes, Proteobacteria* and *Ascomycota, Basidiomycota* for bacteriome and mycobiome, respectively. *Fusobacteria* were detected in samples of both cohorts.

**Figure 3.**
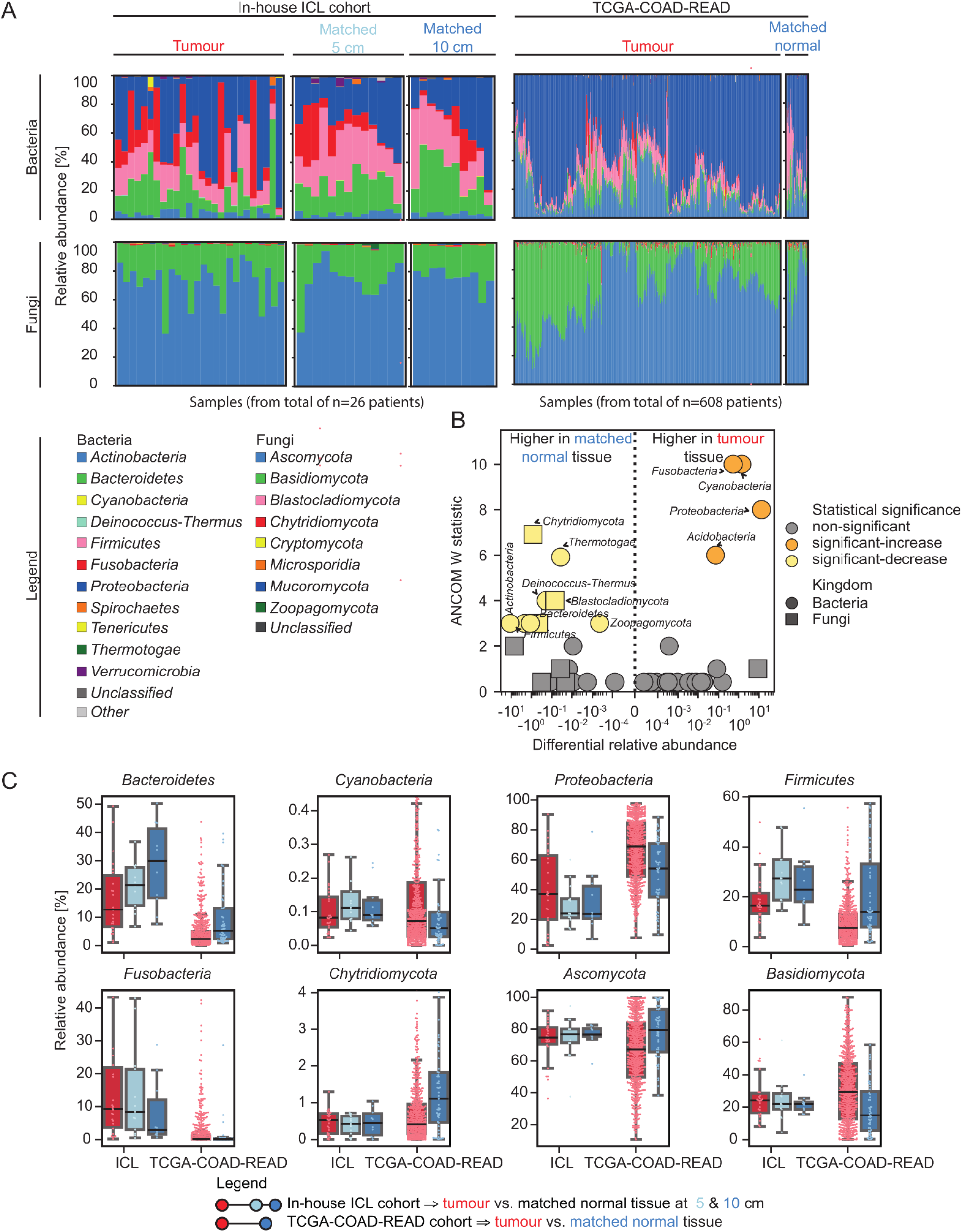
Distinct compositional differences between bacterial and fungal between on-tumour and off-tumour adjacent mucosa from CRC patients. **A.** Bacterial and fungal composition (relative abundance, in percentage) at the phylum taxonomic level from tumour and adjacent normal mucosa from the in-house ICL and TCGA-COAD-READ cohorts. **B.** Volcano plot depicting the relationship between statistical significance using the ANCOM W as metric and differential relative abundance in on-tumour compared to off-tumour adjacent mucosa of bacterial and fungal phyla from samples of the TCGA-COAD-READ cohort. **C.** Boxplot with overlaid swarmplot depicting the distribution of key bacterial and fungal phyla identified as differentially abundant from analysis in **B** in on-tumour and off-tumour adjacent mucosa from the in-house ICL and TCGA-COAD-READ cohorts.

We systematically investigated differences in the bacteriome and mycobiome phyla in tumour and paired matched normal tissue using the TCGA-COAD-READ cohort as discovery cohort due to its larger sample size (**Fig. 3B**). The bacteriome of CRC tumours was enriched with *Fusobacteria, Cyanobacteria, Proteobacteria* and *Acidobacteria* and depleted of *Actinobacteria, Bacteroidetes, Deinococcus-Thermus, Firmicutes* (**Fig. 3B**). Tumour tissue was enriched with fungi such as *Basidiomycota* and depleted of *Ascomycota*. We confirmed these findings in the in-house ICL cohort (**Fig. 3C, Sup. Fig. 2**), where we additionally observed an increase in RA of *Bacteroidetes* and *Ascomycota* with the distance from the tumour. In contrast, *Proteobacteria* and *Firmicutes* RA drastically differ when comparing tumour with matched tissue, regardless of distance.

Next, we investigated differences between microbiome composition in tumour and matched normal tissue at higher taxonomic resolution (**Fig. 4, Sup. Fig. 3**). We visualised the average differences in the bacteriome (**Fig. 4A**) and mycobiome (**Fig. 4B**) from phylum to species taxonomic ranks in the TCGA-COAD-READ cohort. **Fig. 4C, D** displays bacteria (**Fig. 4C**) and fungi (**Fig. 4D**) RA from phylum to family taxonomic rank for phyla identified as statistically significant and with an average difference above 20% when comparing tumour with normal tissue. Tumour tissue was enriched with *Proteobacteria,* largely from the *Gammaproteobacteria* class from the *Enterobacteriales* family; *Fusobacteria,* largely from the *Fusobacteriaceae* family [1,10–12]. Matched normal tissue was enriched with *Firmicutes,* largely from the *Bacillales/Lactobacillales* and *Clostridiales* families, and *Bacteroidetes* from the *Bacteroidaceae* family. We observed a shift in the mycobiome with tumours exhibiting higher *Ascomycota* from the *Saccharomycetes* family, and an overall decrease in *Basidiomycota* stemming from a marked reduction in the *Agaricomycetes* family and a modest increase in *Ustilaginomycotina,* primarily from the *Malasseziales* family.

**Figure 4.**
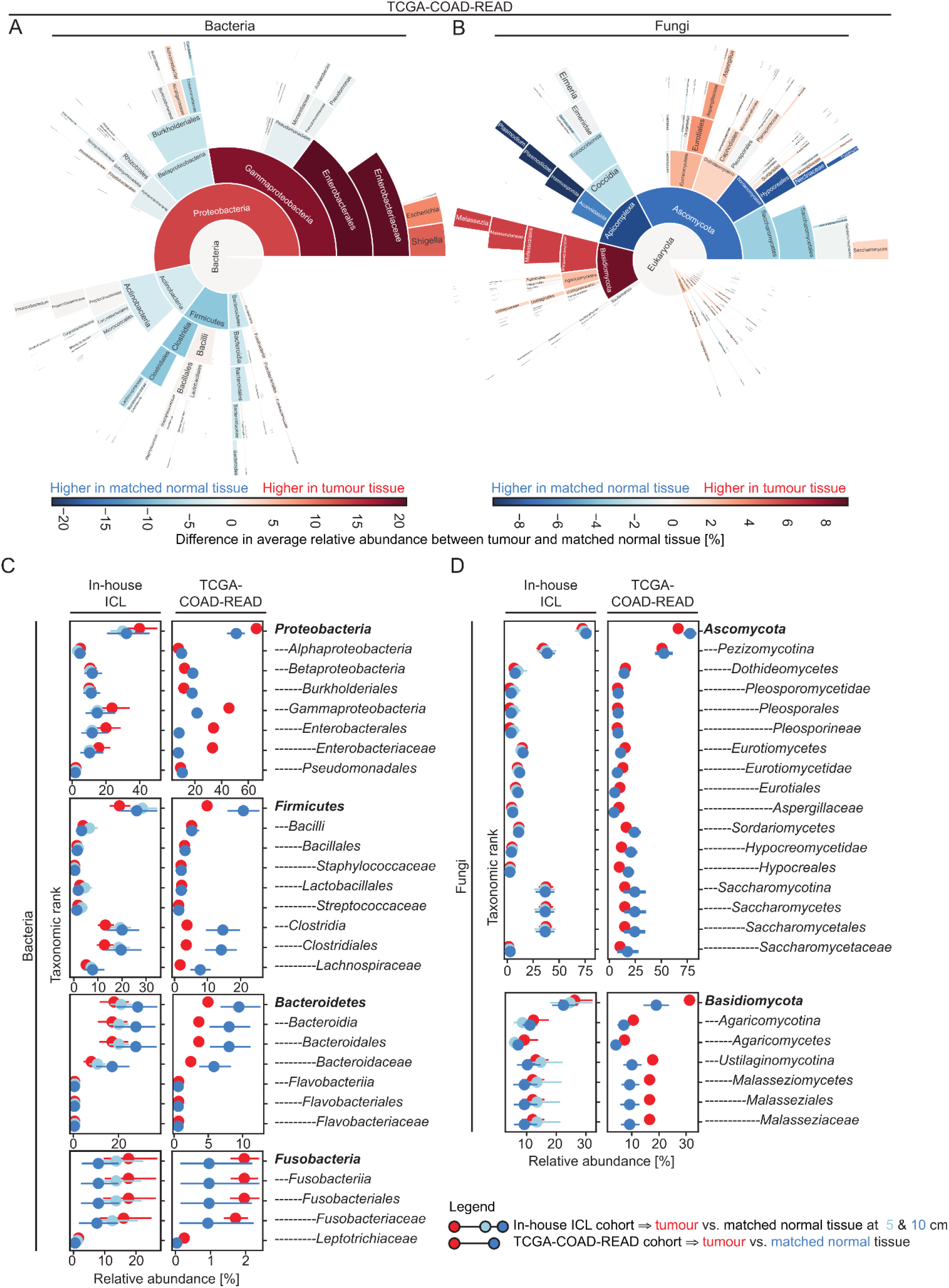
Distinct compositional differences in the bacteriome and mycobiome at higher taxonomic resolution between on-tumour and off-tumour adjacent mucosa from CRC patients. **A-B.** Sunburst visualisation highlighting the average difference in relative abundance of the bacteriome (**A**) and mycobiome (**B**) between on-tumour and off-tumour adjacent mucosa in patients of the TCGA-COAD-READ cohort. Each sunburst circle encodes a different taxonomic rank from (sub)-kingdom (innermost circle) to phylum, class, order, family to genus (outermost circle). **C-D.** Relative abundance of bacteria (**C**) and fungi (**D**) from phylum to family taxonomic rank for phyla identified as statistically significant and with an average difference above 20% when comparing tumour with adjacent normal mucosa for patients of the in-house ICL and TCGA-COAD-READ cohorts.

### Identification of four bacteriome- and mycobiome-derived subtypes

We applied unsupervised agglomerative consensus clustering in TCGA-COAD-READ tumours on RA of bacterial and fungal families and identified n=4 robust microbial-based patient clusters, termed C1 (n=296, 49%), C2 (n=208, 34%), C3 (n=59, 10%) and C4. (n=45, 7%) (**Fig. 5A**). Annotation of the average RA of bacterial and fungal families by patient cluster in **Fig. 5A.2** highlights key drivers of microbial assignments. A distinguishing feature of patients classified as microbial subtype C2 is the higher RA (average 69.9% in C2 vs. 12.9% to 19.0% in C1,C3-C4) of *Enterobacteriaceae,* a resident intestinal microbe implicated in CRC initiation and progression [3,35,36]. Patients classified as microbial subtype C3 featured a higher RA of *Saccharomycetaceae* (average 88.4% in C3 vs. 1.2% to 2.2% in C1-C2,C4). Patients classified as microbial subtype C4 featured a higher RA of *Aspergillaceae* (average 49.5% in C4 vs. 0.4% to 10.4% in C1-C3). A complete account of the bacterial and fungal families identified as differentially abundant across microbial subtypes is reported in **Sup. Table 4**. Next, we set out to subtype the remaining samples included in our analysis. Thus, we built a classification decision tree using as input the composition of bacterial and fungal families from the TCGA-COAD-READ cohort using the microbial subtyping assignments as target to predict. The decision tree (**Fig. 5B**) was able to recapitulate with high accuracy (~99%) the microbial subtyping assignments from the TCGA-COAD-READ cohort.

**Figure 5.**
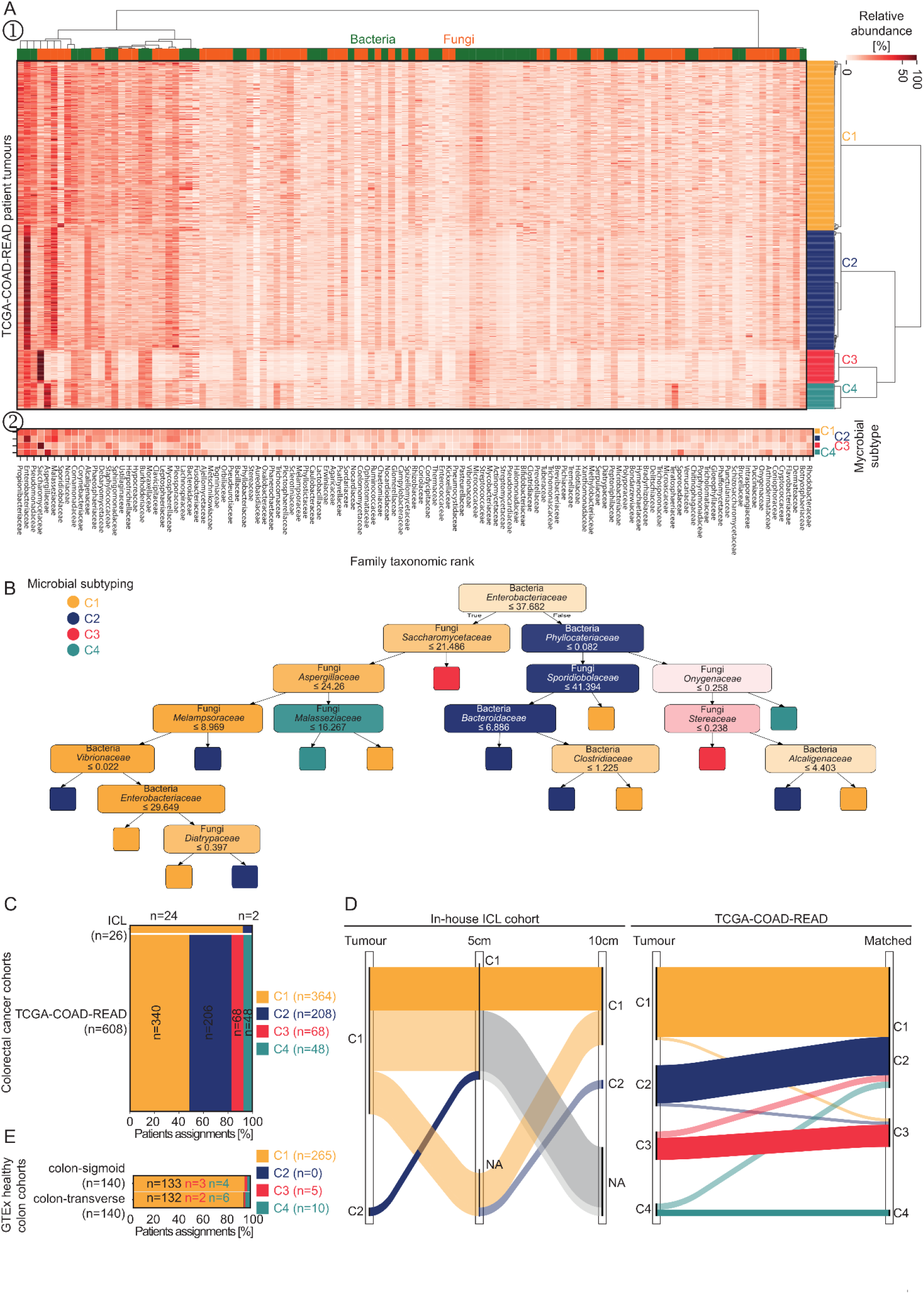
Consensus clustering of the bacteriome and mycobiome classified CRC patients into 4 microbial-based subtypes. **A.** Unsupervised agglomerative consensus clustering of relative abundance of bacterial and fungal families applied to primary tumours of patients from the TCGA-COAD-READ cohort identifies 4 subtypes of patients (C1-C4). Bottom insert heatmap depicts the average relative abundance by subtype. **B.** Classification decision tree trained on the TCGA-COAD-READ cohort, using as input the composition of bacterial and fungal families and the microbial subtyping assignments as target class, identifies key drivers in microbial clustering assignments and can be used to subtype unseen tissue samples with high accuracy (~99%). **C.** Breakdown of subtype assignments for tumour resections of patients of the in-house ICL and TCGA-COAD-READ cohorts. **D.** Sankey diagram tracking the correspondence between subtype assignment in tumours compared to off-tumour adjacent mucosa for patients of the in-house ICL and TCGA-COAD-REAd cohorts. **E.** Breakdown of subtype assignments for sigmoid and transverse colon tissue sampled from healthy subjects of the GTEx consortium.

We used the decision tree to assign microbial subtypes for the ICL cohort (**Fig. 5C**). The majority of the ICL tumours were classified as C1 (n=24/26, 92%) while the remaining 2 tumours (8%) were assigned to subtype C2. We did not identify tumour samples from the less commonly found subtype C3 and C4, attributable to the small sample size of the cohort. While we developed the microbial subtyping for colorectal tumours, we sought to investigate the relationship between microbial signatures in the tumour and paired matched adjacent local and distant mucosa. The vast majority of specimens sampled from matched adjacent mucosa was classified as C1 in both the ICL (n=22/23, 96%) and TCGA-COAD-READ (n=40/51, 78%) cohorts (**Fig. 5D**). When comparing the microbial subtyping of paired tumour-matched normal samples, we observed a consistent and robust shift towards the C1 subtype in adjacent mucosa in both cohorts. All 5 patients with a complete set of samples from tumour and local and distant matched mucosa were classified as C1 in the ICL cohort. In the TCGA-COAD-READ cohort, 22 out of 23 patients with bulk tumour tissue classified as C1 subtype retained the C1 assignment for the matched adjacent mucosa. Twelve out of 13 adjacent mucosa samples shifted to C1 subtype from TCGA-COAD-READ patients with tumours classified as C2 subtypes. Matched adjacent mucosa from tumours classified as C3 or C4 subtype either retained the same assignment as the corresponding tumour or switched to C2 subtype (**Fig. 5D**). Taken together, these results suggest that the C1 subtype may encode a microbial signature more akin to a healthy colorectal microbiota. To test this hypothesis, we estimated the bacterial and fungal composition and assigned microbial subtypes to samples collected from the sigmoid and transverse region of the colon from n=140 healthy subjects (**Sup. Table 5)**of the Genotype-Tissue Expression (GTEx) consortium (**Fig. 5E**). Indeed, n=265 out of 280 (95%) of the colon samples were classified as C1, supporting the hypothesis that this subtype may recapitulate a ‘healthy’ colon bacteriome and mycobiome.

### Immune and signalling relevance of the microbial subtypes

Research highlighting the interaction between the microbiome and anticancer immunosurveillance prompted us to investigate the association between the identified microbial signatures and host TME.

We computed the composition of cell types, including fibroblasts, endothelial and immune cells using the MCP-counter algorithm [37] in patients tumour samples of the TCGA-COAD-READ cohort (**Fig. 6A**). We identified higher abundance of endothelial cells and fibroblasts when comparing abundance in patient tumours classified as C1 vs. C2. C1 tumours had higher abundance of T cells, CD8 T cells, monocytic lineage and neutrophils compared to C2-tumours. Interestingly, C4-tumours presented the highest average fibroblasts and endothelial cell content accompanied by high T cell, cytotoxic lymphocytes, B lineage and myeloid dendritic compared to C1-C3 tumours. We explored whether regulation of established immunomodulators [38], such as antigen presentation, co-inhibitor/stimulator, ligand, receptor and cell adhesion properties, differed by microbial subtype, (**Fig. 6B**). C3-C4 tumours displayed the lowest and highest average expression across the vast majority of immunomodulators compared to C1-C2 tumours, corroborating cell abundance findings (**Fig. 6A**).

**Figure 6.**
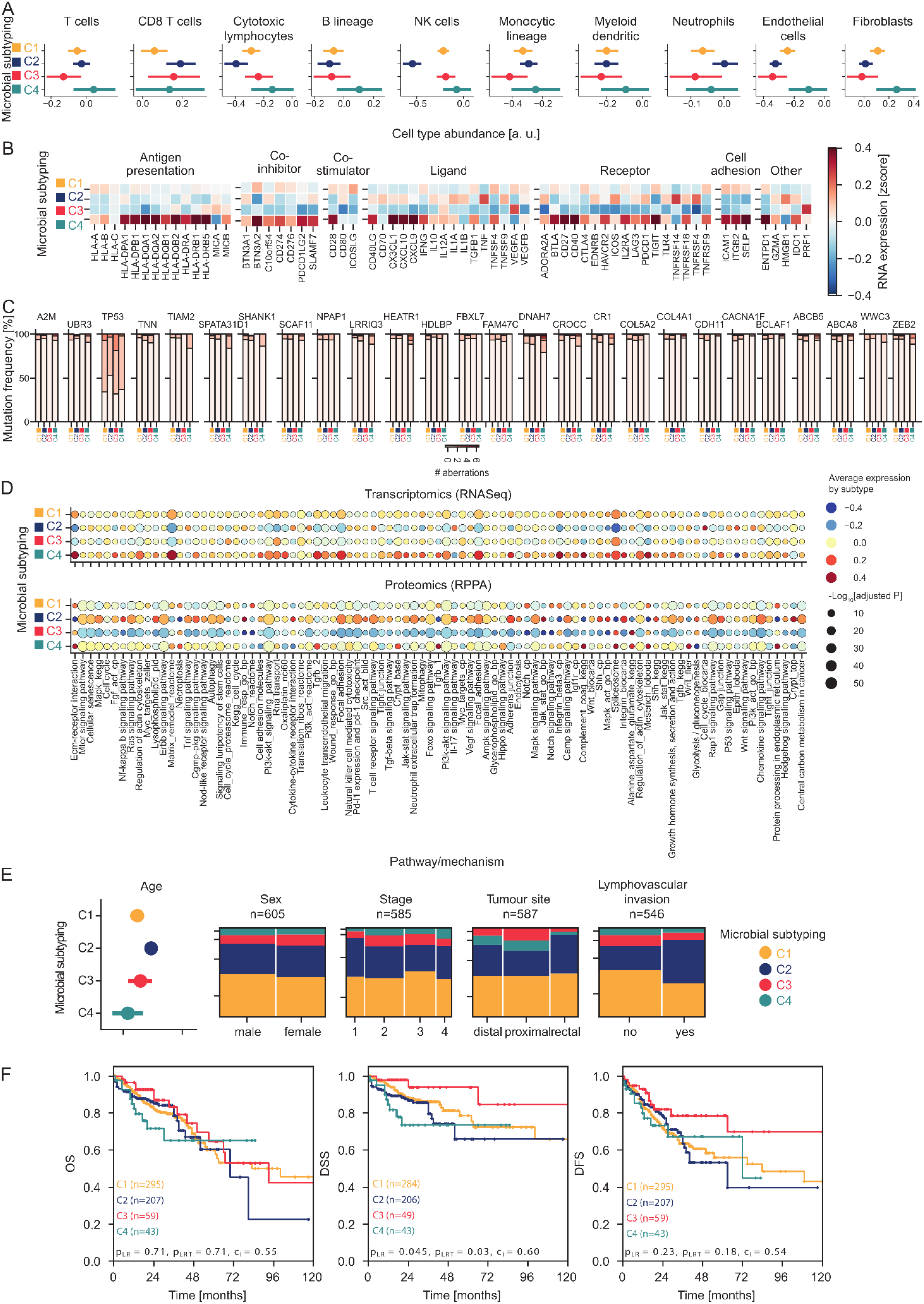
Clinical, molecular and immune underpinning and relevance of microbial subtyping in CRC patients. Analyses were performed in tumours resected from patients of the TCGA-COAD-READ cohort.. **A.** Cell type abundance (average with error bars spanning the 95% CIs) by microbial subtype for fibroblasts, endothelial cells and key immune cells. **B.** Average expression (mean-centred and scaled to unit variance) for key genes involved in immune regulation by microbial subtype. **C**. Breakdown of mutational aberrations by microbial subtype for genes identified as statistically significantly affected by microbial subtype. **D.** Subtype-specific average gene and protein expression signatures for key pathways and gene sets identified as statistically significantly de-regulated by microbial subtype in both transcriptomic- and proteomic-based enrichments analysis. **E.** Association between microbial subtyping and selected clinico-pathological and demographic characteristics of the patients of the TCGA-COAD-READ cohort. **F.** Overall-(OS), disease-free (DSS) and disease-free (DFS) Kaplan-Meier survival curves by microbial subtype.

### Microbial subtypes-specific association with host signalling pathways

We characterised the host biology associated with the microbial signatures by identifying subtype-specific mutations, genes and proteins. The tumour suppressor gene TP53, A2M along with other genes implicated with cancer hallmarks such as epithelial to mesenchymal transition (EMT), focal adhesion, ABC transporters and signalling of WNT-β catenin and IL6-JAK/STAT3 featured among the 27 mutations whose frequency of occurrence differed by microbial subtype (**Fig. 6C, Sup. Table 6**). In line with the immunological findings, C3 tumours presented the lowest number of mutational aberrations, while C4 tumours had the highest.

We performed functional enrichment analysis against a curated list of gene sets encompassing cancer hallmarks and established signatures and cellular processes using as input genes and proteins identified as statistically significantly different by microbial subtypes. Matrix remodelling, spliceosome, mitotic spindle, TNF-α signalling and stromal composition, EMT, WNT-β, p53 and hippo signalling featured among the pathways robustly enriched in analyses performed using both transcriptomic- and proteomic-based profiles (**Fig. 6D**). An increase in cytokine-cytokine receptor interaction and IL-17/SHH signalling along with lower activation of the spliceosome machinery in C2 tumours and matrix remodelling in C2-C3 tumours corroborated cell type and immunoregulation analyses.

### Clinical relevance of the microbial subtypes

The microbial subtypes were associated with clinical phenotype and outcome (**Fig. 6E, Sup. Table 9**). C3-C4 tumours were resected almost exclusively from the colon region, while C1-C2 included ~30% rectal cases. Patients classified as C2 had a history of colon polyps and other malignancies and had lymphovascular invasion and lower prevalence of p53 mutations. C3 tumours had the lowest number of mutations across all microbial subtypes and exhibited longer disease-free survival. C4 patients presented with more advanced T stage tumours, largely from the colon region at younger age compared to the other subtypes. We found no statistically significant association between microbial subtypes and microsatellite instability and neither the consensus molecular subtype (CMS, [39]) nor the cancer-intrinsic subtype (CRIS, [40]), despite having observed such relationships in individual bacteria and fungal families (**Sup. Fig. 4**).

We found statistically significant differences when comparing curves for disease-specific (DSS, logrank p=0.045), but not overall (OS) or disease-free survival (DFS) in the TCGA-COAD-READ cohort (**Fig. 6E**). Univariate Cox regression models highlighted a group of patients (C3) with more favourable outcomes, which were confirmed in multivariate analysis adjusting for clinico-pathological characteristics (**Sup. Fig. 5**).

### Application of microbial subtypes in the pan-cancer settings

Having established the biological and clinical relevance of the identified microbial subtypes in CRC, we investigated whether their applicability extended to the pan-cancer settings.

We profiled the bacteriome and mycobiome and assigned microbial subtypes to primary tumour and matched-adjacent mucosa samples of n=9493 patients from n=28 additional cancer types. The vast majority of tumours were classified as C1 (n=6170/9493, 65%). All 4 microbial subtypes were identified in n=18 (62%) cancer types. Thoracic cancers such as breast (TCGA-BRCA) and lung (adenocarcinoma: TCGA-LUAD; squamous cell carcinoma: TCGA-LUSC) resembled the most the relative frequency of microbial assignments observed in the TCGA-COAD-READ cohort, (**Fig. 7A**). As observed in CRC, the majority of adjacent mucosa was classified as C1 (n=405/728, 56%) and a robust shift towards C1 assignments was observed when comparing subtypes from paired tumour-adjacent mucosa samples, (**Fig. 7B**).

**Figure 7.**
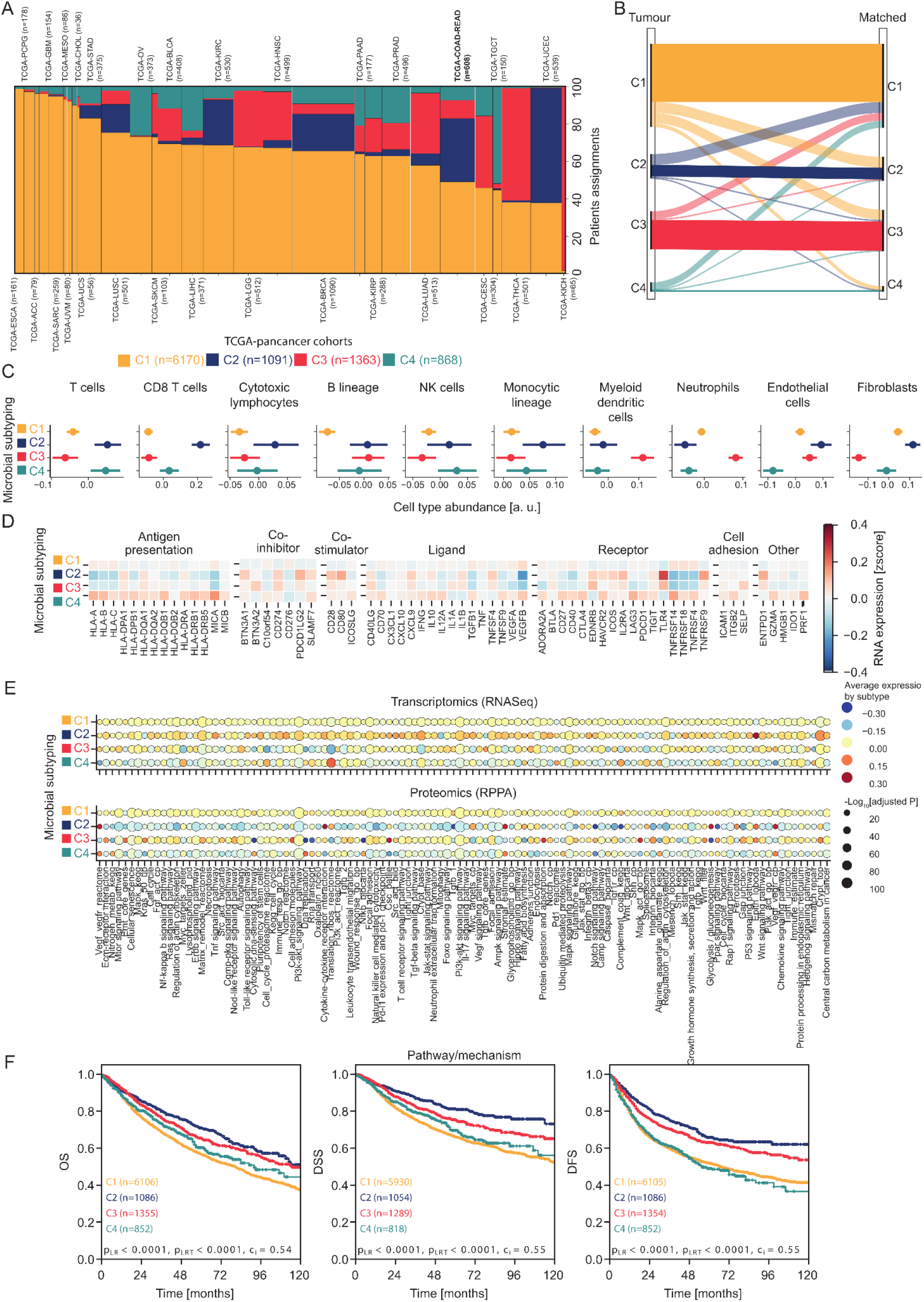
Validation of clinical, molecular and immune underpinning and relevance of microbial subtyping in patients diagnosed with solid pan-cancers. Analyses were performed in solid tumours resected from patients diagnosed with 28 cancer types in addition to CRC from the TCGA pan-cancer collection. **A.** Breakdown of subtype assignments for tumour resections by cancer type. **B.** Sankey diagram tracking the correspondence between subtype assignment in tumours compared to off-tumour adjacent mucosa across all pan-cancer collections. **C.** Cell type abundance (average with error bars spanning the 95% CIs) by microbial subtype for fibroblasts, endothelial cells and key immune cells. **D.** Average expression (mean-centred and scaled to unit variance) for key genes involved in immune regulation by microbial subtype. **E.** Subtype-specific average gene and protein expression signatures for key pathways and gene sets identified as statistically significantly de-regulated by microbial subtype in both transcriptomic- and proteomic-based enrichments analysis. **F.** Overall-(OS), disease-free (DSS) and disease-free (DFS) Kaplan-Meier survival curves by microbial subtype.

The immunological and signalling findings in CRC were largely replicated in the pan-cancer settings, confirming the involvement of key cancer and inflammation signalling pathways including cytokine-cytokine interactions, toll-like receptor signalling, matrix remodelling, focal adhesion, EMT, WNT-β and angiogenesis (**Fig. 7C-E, Sup. Tables 10-11**). C2 and C4 tumours exhibited higher abundance of T cells, CD8 T cells, cytotoxic lymphocytes, monocytic lineage compared to C1 tumours (**Fig. 7C**). C4 tumours presented the highest average expression of immunomodulators, particularly in terms of antigen presentation (**Fig. 7D**).

We observed strong differences in clinical outcome in the TCGA pan-cancer settings, with C2-C3 tumours exhibiting longer overall-, disease-free- and -recurrence-free-survival compared to tumours classified as C1,C4 (**Fig. 7F**). Cox regression models by cancer type demonstrated that the stratification potential of the identified microbial subtypes may extend beyond CRC (**Sup. Table 12**).

## Conclusions

Alterations in the gut microbiome have been linked to CRC development, progression and therapy response [1–5,10–12]. We systematically characterised the bacteriome and mycobiome of on- and off-tumour adjacent mucosa samples from two richly-annotated CRC cohorts to delve into the role played by the microbiome in mediating oncogenesis and uncover host-characteristics that may be amenable to microbial-based therapeutics. Furthermore, in our in-house cohort, we collected tumour resections at local and distant sites from the tumour margins, allowing us to characterise the microbiome in spatially distinct peri-tumour regions.

On-tumour tissue was enriched with *Fusobacteria*, *Proteobacteria*, *Basidyomicota* and depleted of *Bacteroidetes*, *Firmicutes*, *Ascomycota* compared with adjacent off-tumour mucosa. Abundance of *Fusobacteria*, largely from the *Fusobacteriaceae* family and *Fusobacterium nucleatum* species, have been implicated in the aetiology [12,31,41], clinical phenotype [11,33,34,42] such as tumour location, microsatellite instability, assignment to transcriptome-based molecular subtyping, clinico-pathological features, therapy response [32,43] and, ultimately, clinical outcome [11,33,44]. This study identified a depletion of *Firmicutes,* driven by *Clostridia,* but not *Bacilli,* and *Bacteroidetes,* driven by *Bacteroidia,* but not *Flavobacteriia,* corroborating findings in the CRC settings [45]. The depletion of these bacteria, which ferment dietary fibres into butyrate, may result in impairment of structural and immune homeostasis of the gut. Higher *Gammaproteobacteria (Proteobacteria)* RA on-tumour compared to off-tumour adjacent mucosa corroborates previous findings in CRC, pancreatic and non-small cell lung cancer. Boesch *et al.* reported lower PD-L1 expression and poorer response to checkpoint immunotherapy in non-small cell lung cancer patients with higher abundance of *Gammaproteobacteria* [18]. The gut mycobiome has received limited attention compared to the bacteriome because of their relatively lower abundance and difficulty in culturing. In line with findings from Coker *et al.* [14], we observed no differences in neither within-sample nor across-samples α- and β-diversity measures when comparing the mycobiome of on-tumour to off-tumour adjacent mucosa samples. Nevertheless, we observed a robust shift in fungal composition, namely an enrichment of *Basidiomycota* and depletion of *Ascomycota*, when transitioning from on-tumour to off-tumour adjacent mucosa. A higher *Basidiomycota:Ascomycota* ratio has been reported as indicative of fungal dysbiosis in CRC [14–16], other cancers [46] and autoimmune diseases [47–49]. The higher *Basidiomycota* RA observed on-tumour stemmed from *Malasseziaceae*, which has been linked to CRC, other gastrointestinal cancers [46,50], multiple inflammatory [51,52] and skin disorders [53].

Recent findings underscore that CRC tumorigenesis may be promoted *via* alterations of the microbiome ecosystem, rather than by the infection of specific drivers [1], suggesting that microbial compositional signatures rather than individual pathogens may serve as more appropriate readout of a tumour state. Furthermore, it has become increasingly recognised that the oncogenic impact of the same microbial signature may be dampened or exacerbated by the interaction with the unique signalling biology and TME of the host tumour. These observations prompted us to develop a microbial subtyping framework to identify distinct patient’s groups based on the compositional signatures of their bacteriome and mycobiome. C1 tumours were enriched for *Propionibacteriaceae*, a symbiont bacterial family with probiotic properties [54]. C2 tumours boasted a high RA of *Enterobacteriaceae* and *Malasseziaceae,* implicated in cancer initiation and progression [3,36,46,50], but had lower abundance of *Fusobacteriaceae, Bacteroidaceae* and *Ruminococcaceae.* C3 tumours featured a higher RA of *Saccharomycetaceae*, which has been linked to cachexia, inflammation and leaky gut mucosal barrier [55] while *Aspergillaceae* identified C4 tumours. The microbial subtypes were associated with distinct prognosis and clinical patient phenotypes such as staging, tumour location, history, lymphovascular invasion and TP53 status. Interestingly, we did not observe an association between the microbial subtypes with microsatellite instability, CMS-[39] and CRIS-[40] subtypes, despite observing such a relationship with individual families, particularly *Bacteroidaceae* and *Fusobacteriaceae,* as previously reported [11,33,42]. These findings further underscore the orthogonal value of microbial fingerprinting in addition to transcriptomic-based subtyping and more conventional stratification signatures and clinical markers. The critical role in oncogenesis and therapy response played by the microbiome may extend to a broader range of cancer types, in addition to CRC [6,7]. Breast cancer, lung adenocarcinoma and lung squamous cell carcinoma resembled the relative frequency of microbial assignments observed in the CRC settings. Our findings are consistent with the notion that the microbiome of tumours reflects a mixture of site-specific microflora [6,7] coupled with potentially pathogenic bacteria and fungi that may share common characteristics across cancer types. Bacteria and fungi considered pathogenic such as *Fusobacterium* and *Malassezia* have been implicated in breast, pancreatic and lung cancers [50,56]. Furthermore, the microbial subtyping demonstrated stratification value in the pan-cancer settings beyond merely representing differences in survival by cancer type, providing the strongest stratification in liver cancer (TCGA-LIHC). In CRC and pan-cancer cohorts, the majority of matched adjacent mucosa samples were classified as C1 and paired tumour-matched normal samples demonstrated a robust shift towards the C1 subtype in off-tumour tissue. Our hypothesis that the C1 subtype may recapitulate a healthier-like microbiome was supported by the fact that microbial subtyping of colon samples from GTEx healthy subjects were categorised as C1 almost exclusively.

Taken together these findings provide possible avenues to design microbiota modulation therapy tailored to specific patients subgroups with distinct bacteriomes and mycobiomes.

## Materials and methods

Detailed methods are provided in the online supplemental materials and methods.

### Patient and public involvement statement

Patients or the public were not involved in the design, recruitment, conduct, reporting and dissemination of this research.

### Data availability

The raw sequencing data for the in-house ICL cohort have been deposited at Gene Expression Omnibus (GEO) with accession number GSE213800, which will be made publicly available upon publication. Processing and analysis code along with bacterial and fungal estimates included in the study for both the in-house ICL and TCGA pan-cancer collections with corresponding clinical and molecular datasets will be made publicly available and archived upon publication at Zenodo (https://doi.org/10.5281/zenodo.6246345).

## Supporting information

Supplementary figures with captions

Captions for supplementary tables

## Acknowledgements

We wish to thank the patients who kindly donated their samples and made this study possible. This work was funded by a US-Northern Ireland-Ireland Tripartite grant from Science Foundation Ireland and the Health Research Board to JHMP (16/US/3301). The results included here are in part based upon data generated by the TCGA Research Network: https://www.cancer.gov/tcga. The Genotype-Tissue Expression (GTEx) Project was supported by the Common Fund of the Office of the Director of the National Institutes of Health, and by NCI, NHGRI, NHLBI, NIDA, NIMH, and NINDS. The data used for the analyses described in this manuscript were obtained from the GTEx Portal and dbGaP accession number phs000424.v8.p2 on 2020-05-18. We wish to acknowledge the Information Technology department at the Royal College of Surgeons in Ireland and the DJEI/DES/SFI/HEA Irish Centre for High-End Computing (ICHEC) for the provision of computational facilities and support.

## Author Contributions

MS, JK and JHMP conceptualised and designed the study. All authors were involved in collection, preparation, interpretation, validation and critical review of the data. MS and LP prepared and validated the clinical data for the patients of the in-house ICL cohort. SC and ROB prepared the patient samples of the in-house ICL cohort for RNA sequencing. MS performed formal analysis including bioinformatics and statistical analyses. MS created the manuscript figures and supplementary materials. MS and JHMP drafted the manuscript. All authors edited, reviewed, revised and approved the manuscript text. JK and JHMP acquired funding for the study.

