## Supplementary materials and methods for "Identification and characterization of four bacteriome- and mycobiome-derived subtypes in tumour and adjacent mucosa tissue of colorectal cancer patients"

**Materials and Methods** **2**
Clinical cohorts 2
Cancer patients datasets 3
Imperial College cohort (ICL) 3
Colorectal and extended solid pan-cancer collection from The Cancer Genome Atlas (TCGA) 3
Datasets of healthy individuals. 3
Colon samples from the Genotype-Tissue Expression (GTEx) collection. 3
RNA extraction and sequencing for the samples of the Imperial College cohort (ICL) 4
Patients' classification into transcriptomic-based molecular subtypes 4
Estimation of bacterial and fungal composition 6
Microbial subtyping 7
Systematic analysis of the associations between host microbial subtyping and clinical, pathological and molecular features. 8
Characterization of the tumour microenvironment 8
Mutational status 9
Aberrations in gene expression and protein profiles 9
Statistical analysis. 10
Comparative analysis. 10
Differential microbial abundance analysis. 10
Association between microbial subtyping assignments and bacterial and fungal family relative abundance 10
Outcome analysis. 10
Software and libraries 11
References 13

### Materials and Methods

#### Clinical cohorts

This study includes in-house and public datasets profiling the microbiome of n=9730 subjects, spanning patients diagnosed with cancer (n=9518) of the colon and rectum (in-house Imperial College Cohort ICL: n=26; and public TCGA-COAD-READ cohort: n=608) and other solid cancers (TCGA pan-cancer collection, n=28 cancer types in addition to CRC, totalling n=8884) and colon samples from healthy individuals (GTEX collection: n=212). Additionally, matched adjacent tissue was available for n=746 cancer patients (out of n=9553 patients with either tumour and/or matched mucosa, 8%). A breakdown of the number of samples available per tissue and organ across the datasets is available in **Sup. Table 1**. Clinical, demographic and pathological characteristics of the cancer patients for the colorectal and pan-cancer cohorts and healthy individuals are summarised in **Sup. Tables 2-3** and **Sup. Table 5**, respectively.

#### Cancer patients datasets

##### Imperial College cohort (ICL)

Consecutive colorectal CRC cancer patients undergoing surgical resection were accrued at St. Mary's Hospital (Imperial College London, London, UK; *ICL cohort*) between November 2010 and January 2012. All subjects (n=26) provided informed consent and the study received ethical approval by the institution ethical committee (REC reference numbers 07/H0712/112 and 11/LO/0686). Clinical, demographic and pathological characteristics of the ICL patients are reported in **Sup. Table 2**. Fresh frozen tumour samples and matched normal adjacent mucosa tissue resected at 5 and 10 cm from the tumour margins were collected during surgical resection with curative intent. All samples (n=49, **Sup. Table 1**) were assessed by a pathologist who confirmed the presence of tumour and matched normal adjacent mucosa tissue and stored at -80°C. Bacterial profiling by 16S rRNA was performed on samples collected from n=18 patients, as previously described [1].

##### Colorectal and extended solid pan-cancer collection from The Cancer Genome Atlas (TCGA)

Patients diagnosed with a solid cancer accrued by The Cancer Genome Atlas (TCGA network) were considered for inclusion in the study. For CRC-related analysis, we considered for inclusion stage I-IV patients diagnosed with cancer of the colon (COAD) or rectum (READ), referred to as the TCGA-COAD-READ cohort. We extended the analysis to n=28 additional solid cancers from the TCGA collection. In line with what previously described [2], in downstream CRC and pan-cancer analyses, we included only tumour and adjacent mucosa samples with at least a high quality RNASeq experiment from which bacterial and fungal composition could be estimated. Clinical, demographic and pathological characteristics of the TCGA-COAD-READ patients are reported in **Sup. Table 2**.

#### Datasets of healthy individuals.

##### Colon samples from the Genotype-Tissue Expression (GTEx) collection.

We included tissue samples collected post-mortem from the colon-sigmoid (n=140) and colon-transverse (n=144) collected from n=212 healthy individuals from the Genotype-Tissue Expression (GTEx) project [3,4] with available paired-end RNA sequencing experiments (available in dbGaP in SRA format) of good quality (*i.e.* marked for inclusion in the GTEx v8 data freeze). Clinical, demographic and pathological characteristics of the healthy patients with colon-sigmoid or colon-transverse samples from the GTEx cohorts are reported in **Sup. Table 5**.

**RNA extraction and sequencing for the samples of the Imperial College cohort (ICL)**

RNA was extracted from n=49 fresh frozen tissue samples collected from n=26 patients using a standard trizol extraction protocol, as previously described [5]. The resulting RNA pellets were let to air-dry prior to being resuspended in RNase-free water. Quantity and quality of the RNA samples were evaluated using the NanoDrop spectrophotometer (NanoDrop 2000, Thermo Fisher Scientific, MA, USA). Sequencing libraries were prepared by converting the mRNA to cDNA followed by adapter ligation and enrichment of exon-coding sequences by PCR using sequence-specific probes. Using the KAPA-stranded mRNAseq Kit (Illumina), as per manufacturer's instructions. Whole-transcript RNA sequencing was performed with Illumina HiSeq 4000 using a flow cell generating $1 \times 51$ bp reads at VIB Center for Cancer Biology (KU Leuven, Belgium), as previously described [5]. Sequencing data were processed using an existing in-house bioinformatics pipeline consisting of: i) removal of the optical duplicates and sequencing adapters; ii) mapping of the reads to the human reference genome using TopHat-Bowtie; iii) generation of gene-count matrices with HTSeq-count [6].

The raw sequencing data have been deposited at Gene Expression Omnibus (GEO) with accession number GSE213800, which will be made publicly available upon publication.

**Patients' classification into transcriptomic-based molecular subtypes**

For the ICL cohort, raw counts were normalized using the *R* package *DESeq2* using *rlog* as normalization method and '~ *patient_id* + *distance*' as design formula, including only genes (ENSEMBL identifiers) with non-zero counts in more than 10% of the samples. 'Distance' was defined as 'bulk' (tumour tissue), '5cm' (normal mucosa collected at 5cm from the tumour resection margins) and '10cm' (normal mucosa collected at 10cm from the tumour resection margins). For the TCGA-COAD-READ and TCGA-pancacer cohorts, level 4 batch-corrected and normalised gene expression profiles by RNASeq were retrieved from the TCGA PanCanAtlas data-freeze release (EBPlusPlusAdjustPANCAN_IlluminaHiSeq_RNASeqV2.geneExp.tsv from https://gdc.cancer.gov/about-data/publications/pancanatlas).

Tumour samples from patients of the in-house ICL and TCGA-COAD-READ cohorts were classified according to the *Consensus Molecular Subtype* (CMS, [7]) and *Cancer Intrinsic Subtype* (CRIS, [8]).

Patients were classified into CMS groups using the nearest prediction from the *R* package *CMSclassifier* (https://github.com/Sage-Bionetworks/CMSclassifier, [7]). We used nearest predictions from the single-sample (SS) or random forest (RF) method for the in-house ICL and TCGA-COAD-READ cohort, respectively. For the in-house ICL cohort, we used the *rlog*-normalized gene expression as input to the *CMSclassifier*. As previously described [2], for the TCGA-COAD-READ cohort, we retrieved the RF nearest prediction labels provided by Guinney *et al.* (*cms_labels_public_all.txt* from synapse accession number *syn4978511*, [7]). Furthermore, to maximise the number of patients with CMS assignments, we computed nearest prediction RF labels for the whole TCGA-COAD-READ cohort *de novo* and additionally included the CMS assignments for those patients that had not been subtyped as part of the Guinney *et al.* study. For both cohorts, subtype assignments that either could not be called or mapping to multiple CMS classes were classified as undetermined and, thus, set to NOLBL (short for "no label").

Patients were classified into CRIS groups and labelled as CRIS-A to CRIS-E or NOLBL (if Benjamini-Hochberg–corrected false discovery rate (BH.FDR) exceeded 0.2), as described in Isella *et al.* [8]. For the in-house ICL cohort, we used the *rlog*-normalized gene expression as input to the *CMSclassifier*. As previously described [2], for the TCGA-COAD-READ cohort, we applied the CRIS subtyping to the whole TCGA-COADREAD cohort and as CRIS assignments, we included either the labels provided from the Isella *et al.* publication [8] or the labels we computed *de novo* for patients that had not been subtyped as part of the original study.

**Estimation of bacterial and fungal composition**

Microbiota composition in solid tissue samples was estimated from RNASeq experiments from both in-house and public datasets using a subtractive method implemented by the *PathSeq* pipeline (version 2, *PathSeqPipelineSpark* routine, [9]), powered by the Genome Analysis Toolkit engine (GATK, https://gatk.broadinstitute.org/, [10]) and the Apache Spark framework. As input to the pipeline, we used either level 1 protected BAM files accessed via the GDC Data Portal (https://portal.gdc.cancer.gov/) for the TCGA cohorts or FastQ files converted to uBAM files via the GATK *FastqToSam* routine for the remaining datasets (*ICL* and *GTEX*, cohorts), respectively. Briefly, host reads (*i.e.* human) were filtered out and the remaining unmapped reads were aligned to microbial reads based on reference taxonomies for bacteria, fungi and viruses using a (default) *min-clipped-read-length* of 31. Host and microbe references files were retrieved from the GATK Resource Bundle (ftp://gsapubftp-anonymous@ftp.broadinstitute.org/bundle/pathseq/). We ran the PathSeq pipeline on patient samples that met the criteria above. Next, we restricted the analysis to samples which exceeded 10 million primary reads, resulting in the final high quality samples set for downstream analysis. In downstream analyses, we included only specimens collected from colon mucosa from healthy patients (GTEx) and specimens collected from either the primary tumour or the matched off-tumour mucosa ofor cancer patients (in-house ICL and TCGA-pancancer cohorts). Specimens for the TCGA-pancancer collections extracted from recurrence, metastasis or peripheral blood were excluded from the study.

A breakdown of the samples included in downstream analyses tallied by source (in-house, TCGA-COAD-READ, TCGA-pancancer and GTEx) and sample type (tumour and matched adjacent mucosa) is included in **Sup. Table 1**. In downstream analyses, we collapsed microbial relative abundance from multiple samples of the same patient by tissue type (tumour and adjacent mucosa) by mean to yield a single estimate by patient for the tumour and matched normal, when available. We reported relative abundance for bacteria and fungi at the phylum, order, family, genus and species taxonomic rank as a normalized score expressed as percentage of the total relative abundance of the bacterial or fungal kingdom/sub-kingdom.

**Microbial subtyping**

Tumour samples from the patients of the TCGA-COAD-READ cohort were used as the discovery cohort.

Samples were classified into microbial subtypes based on the relative abundance of bacteria and fungi at the family taxonomic rank. Among bacteria and fungi, 510 families were detected in at least a tumour sample of the TCGA-COAD-READ cohort. We used as input to the microbial subtyping framework, 117 bacterial and fungal families detected with a relative abundance above a cut-off in at least 10% of the TCGA-COAD-READ tumours. The cut-off was determined as the mean of the mean relative abundance across all bacterial and fungal families. The relative abundance of the selected 117 bacterial and fungal families in tumour tissue of the patients of TCGA-COAD-READ cohort served as input to unsupervised consensus clustering (fraction of samples set to 0.8, number of permutations set to 100) using agglomerative clustering as base task. The elbow rule was used to determine the optimal number of clusters.

A classification decision tree using the bacterial and fungal families as input and the microbial assignment as target class was trained on the TCGA-COAD-READ tumours. Next, the learnt decision tree model was applied to subtype samples it was not trained on including the adjacent mucosa samples from the TCGA-COAD-READ cohort, tumour and adjacent mucosa samples of the ICL cohort and healthy colon tissue from the GTEx collection. Furthermore, the learnt decision tree model was applied to subtype tumours and adjacent mucosa samples of the TCGA-pancancer collection.

**Systematic analysis of the associations between host microbial subtyping and clinical, pathological and molecular features.**

We focus the analysis on patients diagnosed by the TCGA network with either colorectal cancer (TCGA-COAD-READ) or any additional solid cancer (n=28 additional cancer types) via the GDC Data Portal (https://portal.gdc.cancer.gov/). In line with what previously described [2], in downstream CRC and pan-cancer analyses, we included only tumour and adjacent mucosa samples with at least a high quality RNASeq experiment from which bacterial and fungal composition could be estimated. In downstream analysis, multiple samples per-patient per-sample type (tumour vs. matched normal mucosa) were aggregated to yield a sample per-patient per-sample type. In analysis including the clinical metadata, the analysis was further restricted to those patients not marked as "Redacted" in the supplementary materials of Liu *et al*. study [11].

As previously described [2], we collected and harmonised clinical [11] and molecular characterizations from the TCGA PanCanAtlas data-freeze release (https://gdc.cancer.gov/about-data/publications/pancanatlas) including mutations (*mc3.v0.2.8.PUBLIC.maf.gz*), and expression profiling (level 4 batch-corrected and normalised) of genes (*EBPlusPlusAdjustPANCAN_IlluminaHiSeq_RNASeqV2.geneExp.tsv*), and proteins (*TCGA-RPPA-pancan-clean.txt*). We supplemented these characterizations with additional metadata extracted from other publications such as microsatellite status as well as computing *de novo* TME signatures.

**Clinical-pathological features**

Patients were classified as microsatellite stable (MSS) or unstable (MSI) using a cut-off of 0.4 applied to the MANTIS score retrieved from the supplementary materials of Bonneville et al. [12]. Association between microbial subtyping and clinical-pathological features were assessed using the $\chi 2$ and the Kruskal tests for categorical and continuous variables, respectively. P-values were not adjusted for multiple comparisons.

**Characterization of the tumour microenvironment**

As previously described [2], cell type composition was computationally deconvoluted from bulk tumour gene expression of the TCGA-COAD-READ and TCGA-pancancer cohorts using the *R* package *Microenvironment Cell Populations-counter* (*MCPcounter*, [13]). *MCPcounter* uses marker genes to estimate the abundance (in arbitrary units) of endothelial cells, fibroblasts and 8 immune cell types including T cells, CD8+ T cells, cytotoxic lymphocytes, B lineage, natural killer (NK) cells, monocytic lineage, myeloid dendritic cells and neutrophils. We applied a quantile-transform (*sklearn.preprocessing.QuantileTransformer* with optimal distribution set to *normal*) on level 4 batch-corrected and normalised gene expression profiles by RNASeq retrieved from the TCGA PanCanAtlas data-freeze release (*EBPlusPlusAdjustPANCAN_IlluminaHiSeq_RNASeqV2.geneExp.tsv*) followed by robust scaling (*sklearn.preprocessing.RobustScaler*) prior to applying the *MCPcounter* algorithm, as previously described [2].

Additionally, we computed and reported average expression (mean centred and scaled to unit variance) for key genes involved in different aspects of immune regulation by microbial subtyping. Gene groups were based on the work by Thorssson *et al*. [14] and included *antigen presentation*, *cell adhesion*, *co-inhibitor*, *co-stimulator*, *ligand*, *receptor* and *other*.

**Mutational status**

Somatic mutation data in Mutation Annotation Format (*MAF, mc3.v0.2.8.PUBLIC.maf.gz*) were retrieved from the TCGA PanCanAtlas data-freeze release (https://gdc.cancer.gov/aboutdata/publications/pancanatlas) and restricted to the subset of patients either diagnosed with colorectal cancer (TCGA-COAD-READ) or with other solid cancers (TCGA-pancancer).

As previously described [2], for each patient and each gene, we extracted from the MAF file the number of detected mutational aberrations. As aberrations, we included frame shift deletions and insertions, in frame deletions and insertions, missense and nonsense mutations and splice sites and we excluded the following variants: 3’ flank, 3’ UTR, 5’ flank, 5’ UTR, Intron, RNA, silent and non-stop mutations.

Association between microbial subtyping and mutational status (number of aberrations) was assessed with $\chi 2$ independence tests. As previously described [2], we restricted the analysis to genes with aberrations in at least 5% of patients (n=818 genes out of 21332, ~4%). We reported unadjusted- and Benjamini-Hochberg FDR-corrected mod-likelihood P-values.

**Aberrations in gene expression and protein profiles**

As previously described [2], we performed a systematic and unbiased analysis to identify aberrations in gene expression and protein profiles by microbial subtypes for patients of the TCGA-COAD-READ and TCGA-pancancer collections.

Gene expression (*EBPlusPlusAdjustPANCAN_IlluminaHiSeq_RNASeqV2.geneExp.tsv*) as level 4 batch-corrected and normalized expression profiles was retrieved from the TCGA PanCanAtlas data-freeze release (https://gdc.cancer.gov/about-data/publications/pancanatlas). We included the 5000 most variant genes for testing the association between gene expression and microbial subtyping.

Protein expression (*TCGA-RPPA-pancan-clean.txt*) as level 4 batch-corrected and normalized expression profiles was retrieved from the TCGA PanCanAtlas data-freeze release (https://gdc.cancer.gov/about-data/publications/pancanatlas). All available proteins (n=189) were included for testing the association between protein expression and microbial subtyping.

Association between microbial subtyping and either gene or protein expression was assessed by Kruskal-Wallis H-tests. We reported unadjusted- and Benjamini-Hochberg FDR-corrected P-values.

Genes and proteins whose expression differed by microbial subtyping were put forward for pathway enrichment analyses carried out with the *python* package *gseapy* which provides a wrapper (function *gseapy.enrichr*) for EnrichR [15], as previously described [2].

**Statistical analysis.**

Statistical significance was set at $P<0.05$ for both comparative and outcome statistical analyses. P-values were adjusted for False Discovery Rate with Benjamini-Hochberg FDR correction (FDR-BH) in systematic analyses. P-values were not adjusted for multiple comparisons in hypothesis-driven or exploratory analyses, as indicated in the captions.

**Comparative analysis.**

Statistically significant differences between groups of patients by clinical, demographic and pathological characteristics reported in **Sup. Tables 2-3, 5, 9** were computed with the *TableOne python* package. The Chi-square test was employed to assess statistical significant differences for categorical variables. The Kruskal test was employed to assess statistical significant differences for continuous variables. Reported P-values were not adjusted for multiple comparison as the variables to assess had been selected *a priori*.

**Differential microbial abundance analysis.**

Differential microbial abundance between tumour and matched adjacent mucosa for bacterial and fungal phyla was computed using the ANCOM method implemented in the the *skbio.stats.composition.ancom* function from *skbio* package. Briefly, the ANCOM method involves computing pairwise log ratios between all the features, here bacterial and fungal phyla, and testing whether there is a significant difference in the phylum ratio with respect to sample type (tumour vs. adjacent mucosa).

**Association between microbial subtyping assignments and bacterial and fungal family relative abundance**

Statistically significant differences in relative abundance of bacteria and fungi at the family taxonomic rank by microbial subtype were assessed by ANOVA tests followed by pairwise post-hoc Tukey tests.

**Outcome analysis.**

As outcome endpoints, we evaluated disease-free (DFS), disease-specific (DSS) and overall (OS) survival where we considered relapse, cancer-related death or death by any cause as an event, respectively. We used Kaplan-Meier estimators and we fit univariate and multivariate Cox proportional hazards regression models to evaluate survival by covariates. We assessed statistical significance with log-rank and likelihood ratio tests, respectively.

For the TCGA-COAD-READ cohort, we fit univariate Cox regression models and 2 multivariate Cox regression models (M1 and M2). Multivariate model M1 included the microbial subtyping and was adjusted by stage. Multivariate model M2 was additionally adjusted by sex, age and tumour site.

For the TCGA-pancancer collection, we fit univariate Cox regression models by microbial subtyping for each individual cancer type. We restricted this analysis to cancer types where at least 2 microbial subgroups were identified with at least 10 patients each.

**Software and libraries**

Download, process and analysis were performed in *python* (version 3.9.5) with custom code using multiple libraries including *pandas, dask, numpy, sklearn, skbio, matplotlib, seaborn, plotly, UpSetPlot, tableone, statsmodels, pingouin, gseapy* and *lifelines*. The complete list of packages and their versions along with a docker image with code and data will be made publicly available and archived upon publication at Zenodo (https://doi.org/10.5281/zenodo.6246345).
