## Supplementary figures with captions for "Identification and characterization of four bacteriome- and mycobiome-derived subtypes in tumour and adjacent mucosa tissue of colorectal cancer patients"

### Supplementary Figure 1

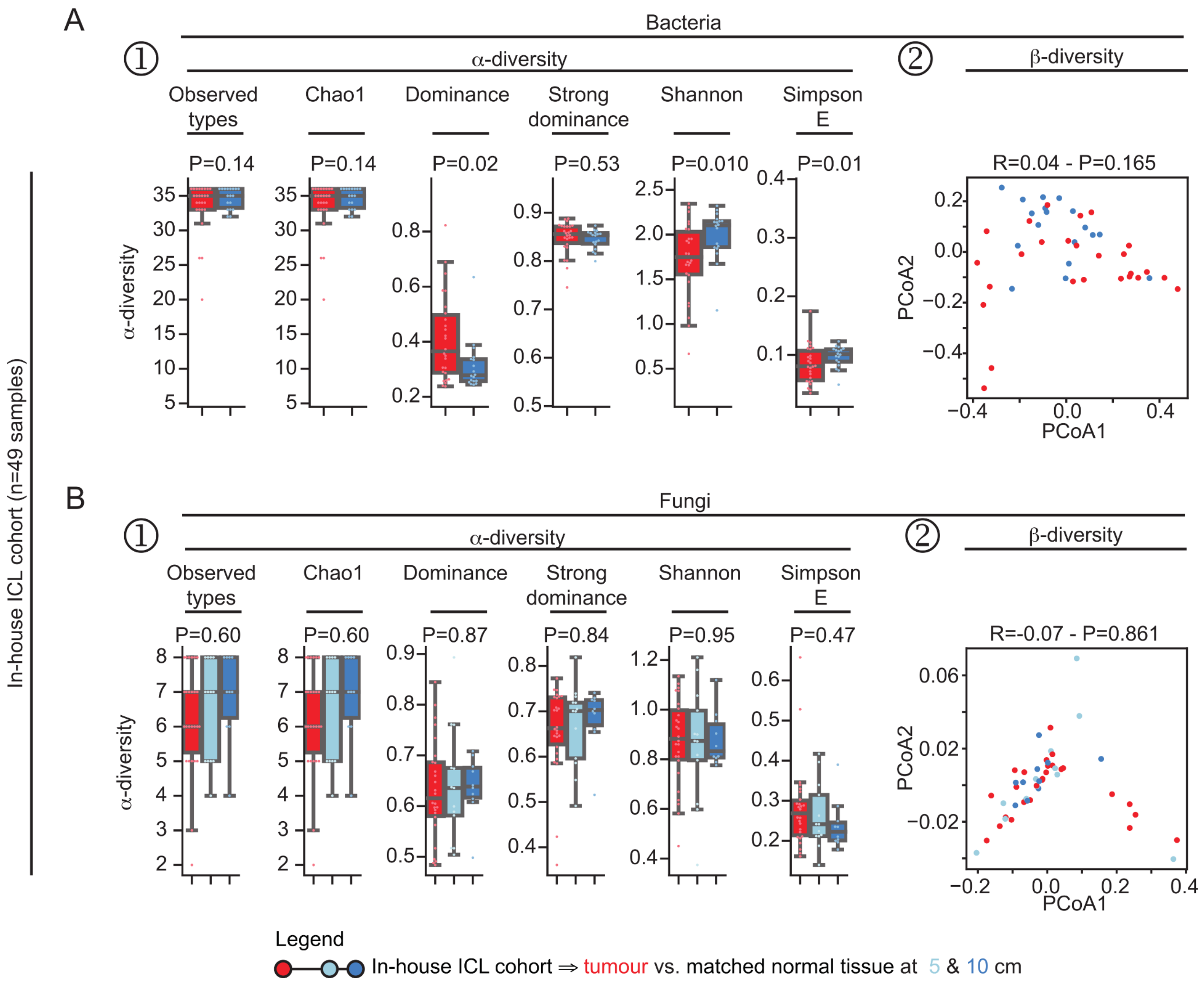


***Supplementary Figure 1. a- and β-diversity metrics for bacterial and fungal phyla in on-tumour tissue samples compared to off-tumour adjacent mucosa in tissue resections from CRC patients of the in-house ICL cohort.***

**A-B.** Within-sample α-diversity (**A.1-B.1**) and across-samples β-diversity (**A.2-B.2**) metrics comparing bacterial (**A**) and fungal (**B**) ecological scores in on-tumour and off-tumour adjacent mucosa samples from patients of the in-house ICL cohort. α-diversity indices included observed types, Chao1, (strong)-dominance, Shannon and Simpson E. β-diversity was quantified by performing unsupervised principal component analysis on Bray-Curtis distances. Statistical significance differences between β-diversity between on-tumour vs. off-tumour adjacent mucosa was performed using analysis of similarities (ANOSIM).

**Supplementary Figure 2**

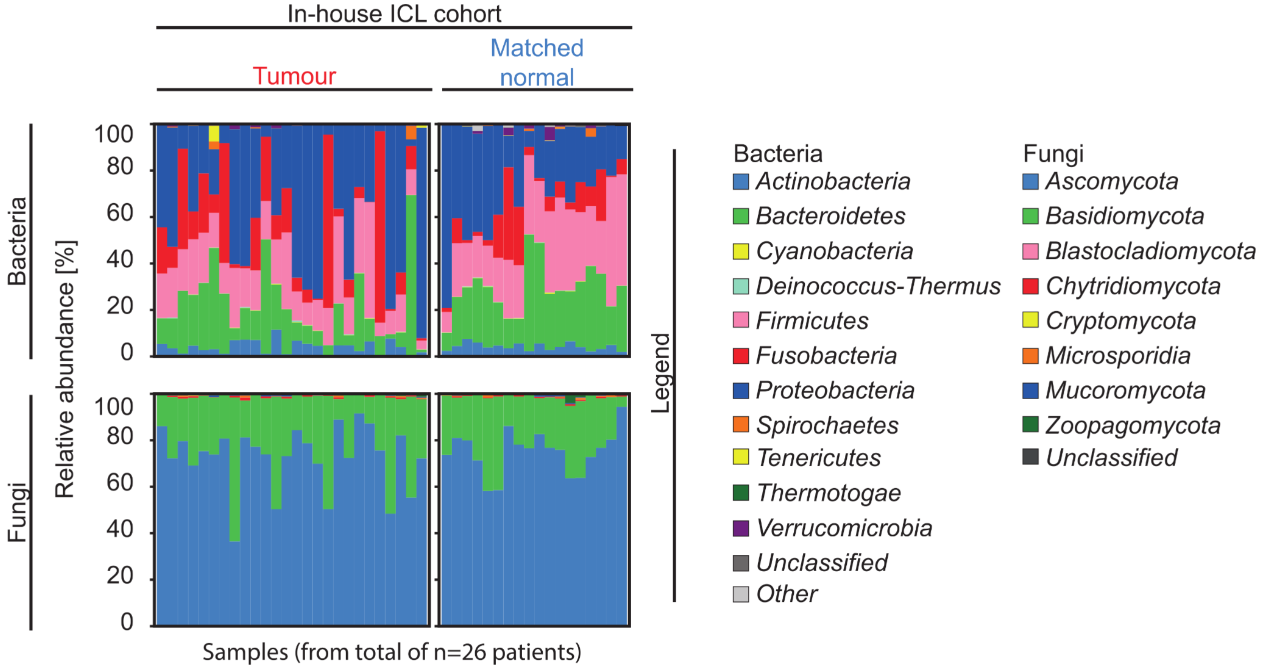


**Supplementary Figure 2.** ***Distinct compositional differences between bacterial and fungal between on-tumour and off-tumour adjacent mucosa from CRC patients of the in-house ICL cohort.***
Bacterial and fungal composition (relative abundance, in percentage) at the phylum taxonomic level from tumour and adjacent normal mucosa from the in-house ICL cohort.

**Supplementary Figure 3**

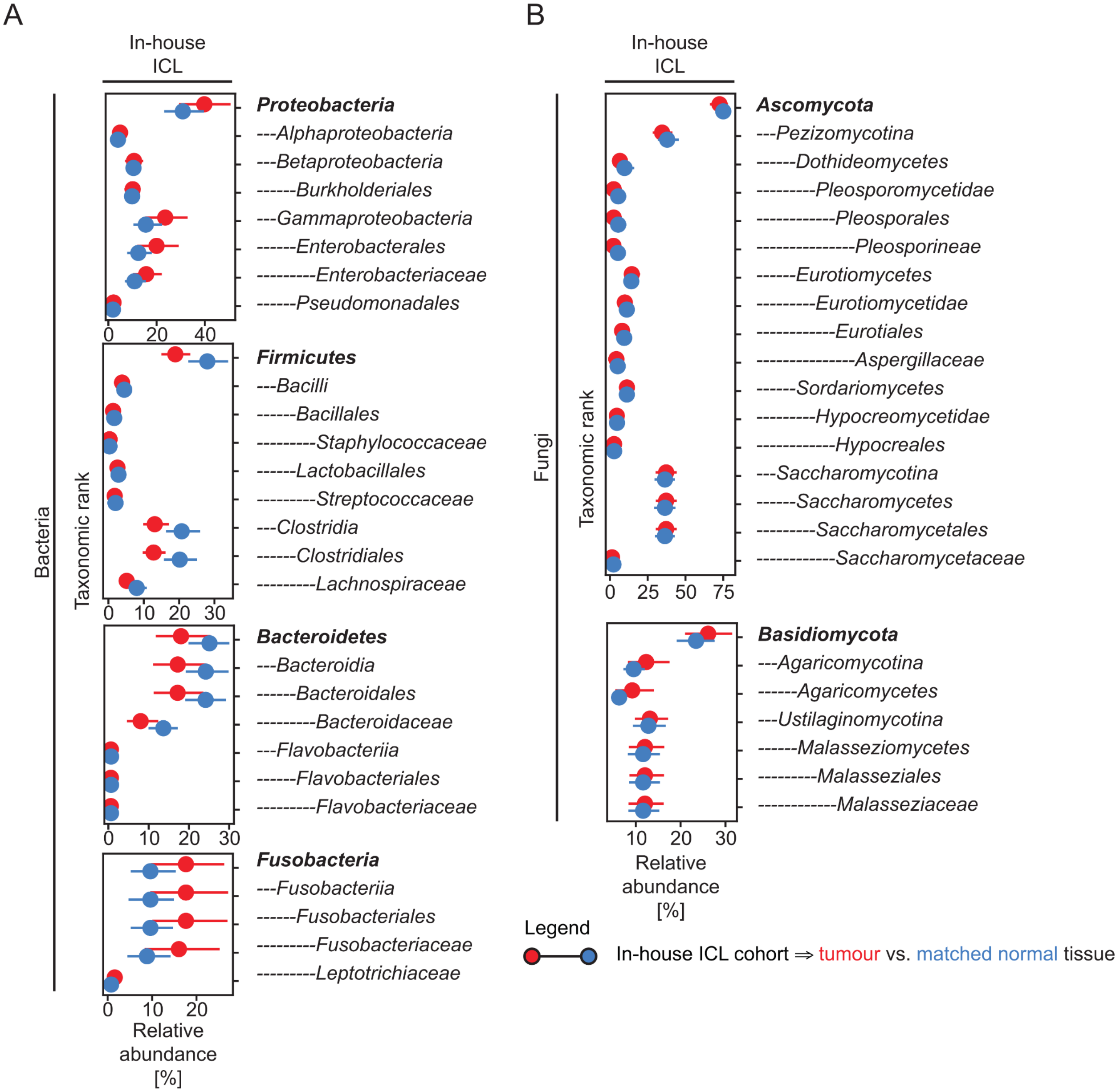


**Supplementary Figure 3.** ***Distinct compositional differences in the bacteriome and mycobiome at higher taxonomic resolution between on-tumour and off-tumour adjacent mucosa from CRC patients of the in-house ICL cohort.***

**A-B.** Relative abundance of bacteria (**A**) and fungi (**B**) from phylum to family taxonomic rank for phyla identified as statistically significant and with an average difference above 20% when comparing tumour with adjacent normal mucosa for patients of the in-house ICL cohort.

**Supplementary Figure 4**

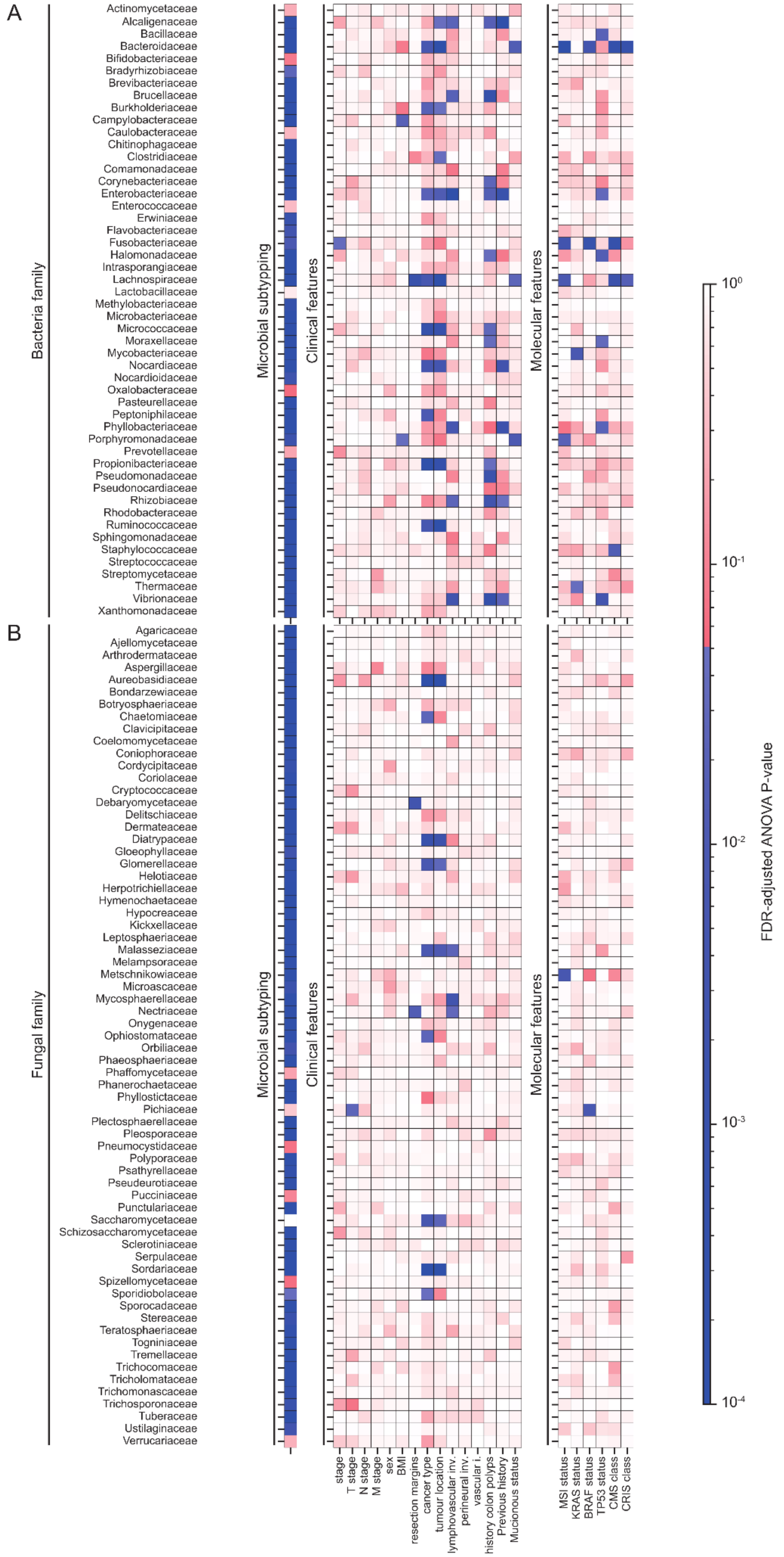

**Supplementary Figure 4.** ***Association between bacteria and fungal families with microbial subtyping and clinical and molecular features in patients of the TCGA-COAD-READ cohort.***

Univariate regression models investigating the association between relative abundance of bacteria and fungal families used as input to the microbial subtyping with clinical and molecular characteristics of the patients of the TCGA-COAD-READ cohort..

**Supplementary Figure 5**

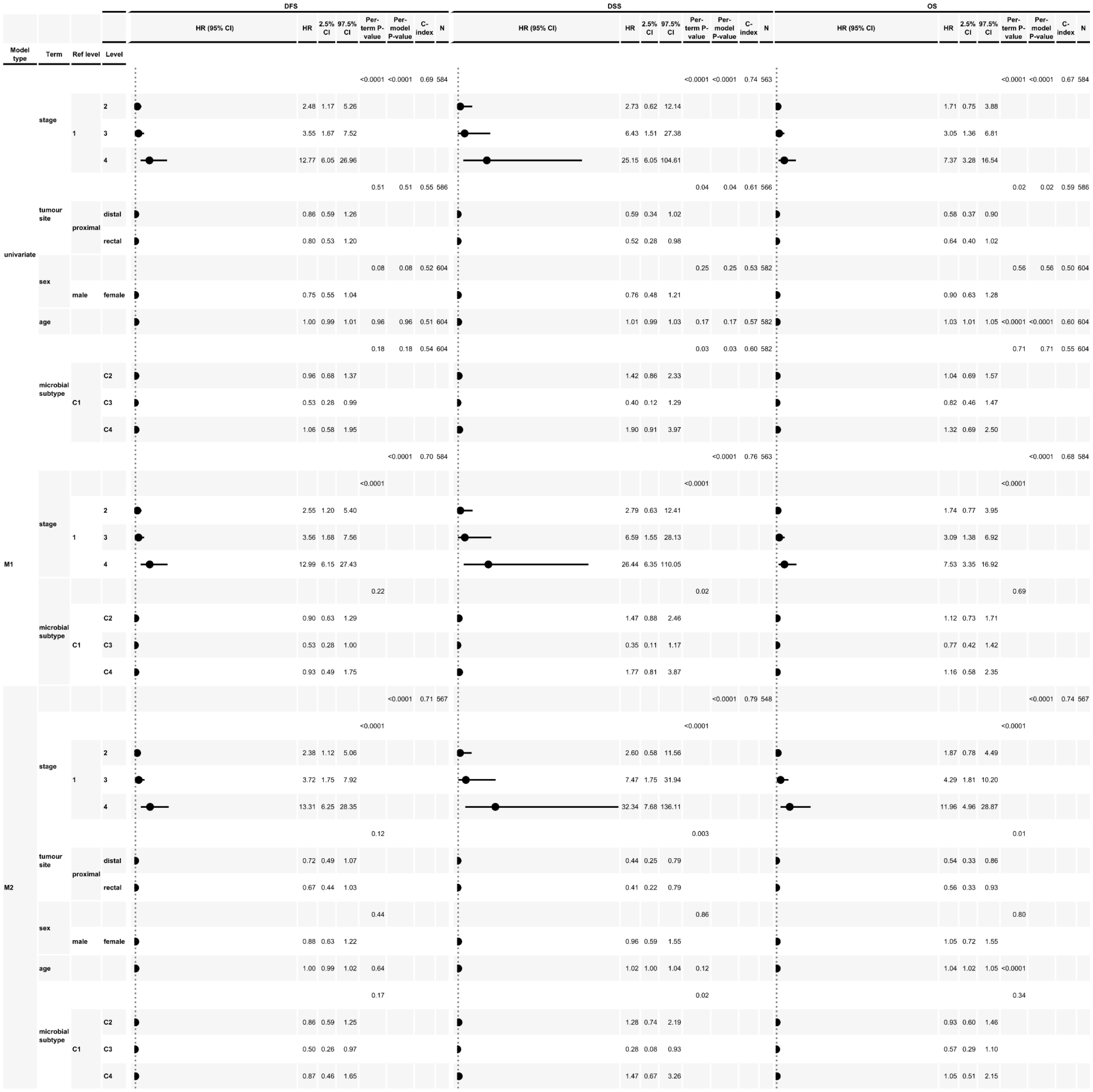

| Model type | Term | Ref level | Level | DFS HR | DFS 2.5% CI | DFS 97.5% CI | DFS Per-term P-value | DFS Per-model P-value | DFS C-index | DFS N | DSS HR | DSS 2.5% CI | DSS 97.5% CI | DSS Per-term P-value | DSS Per-model P-value | DSS C-index | DSS N | OS HR | OS 2.5% CI | OS 97.5% CI | OS Per-term P-value | OS Per-model P-value | OS C-index | OS N |
|---|---|---|---|---|---|---|---|---|---|---|---|---|---|---|---|---|---|---|---|---|---|---|---|---|
| univariate | stage | | | | | | <0.0001 | <0.0001 | 0.69 | 584 | | | | <0.0001 | <0.0001 | 0.74 | 563 | | | | <0.0001 | <0.0001 | 0.67 | 584 |
| | | 1 | 2 | 2.48 | 1.17 | 5.26 | | | | | 2.73 | 0.62 | 12.14 | | | | | 1.71 | 0.75 | 3.88 | | | | |
| | | | 3 | 3.55 | 1.67 | 7.52 | | | | | 6.43 | 1.51 | 27.38 | | | | | 3.05 | 1.36 | 6.81 | | | | |
| | | | 4 | 12.77 | 6.05 | 26.96 | | | | | 25.15 | 6.05 | 104.61 | | | | | 7.37 | 3.28 | 16.54 | | | | |
| | tumour site | | | | | | 0.51 | 0.51 | 0.55 | 586 | | | | 0.04 | 0.04 | 0.61 | 566 | | | | 0.02 | 0.02 | 0.59 | 586 |
| | | proximal | distal | 0.86 | 0.59 | 1.26 | | | | | 0.59 | 0.34 | 1.02 | | | | | 0.58 | 0.37 | 0.90 | | | | |
| | | | rectal | 0.80 | 0.53 | 1.20 | | | | | 0.52 | 0.28 | 0.98 | | | | | 0.64 | 0.40 | 1.02 | | | | |
| | sex | | | | | | 0.08 | 0.08 | 0.52 | 604 | | | | 0.25 | 0.25 | 0.53 | 582 | | | | 0.56 | 0.56 | 0.50 | 604 |
| | | male | female | 0.75 | 0.55 | 1.04 | | | | | 0.76 | 0.48 | 1.21 | | | | | 0.90 | 0.63 | 1.28 | | | | |
| | age | | | 1.00 | 0.99 | 1.01 | 0.96 | 0.96 | 0.51 | 604 | 1.01 | 0.99 | 1.03 | 0.17 | 0.17 | 0.57 | 582 | 1.03 | 1.01 | 1.05 | <0.0001 | <0.0001 | 0.60 | 604 |
| | microbial subtype | | | | | | 0.18 | 0.18 | 0.54 | 604 | | | | 0.03 | 0.03 | 0.60 | 582 | | | | 0.71 | 0.71 | 0.55 | 604 |
| | | C1 | C2 | 0.96 | 0.68 | 1.37 | | | | | 1.42 | 0.86 | 2.33 | | | | | 1.04 | 0.69 | 1.57 | | | | |
| | | | C3 | 0.53 | 0.28 | 0.99 | | | | | 0.40 | 0.12 | 1.29 | | | | | 0.82 | 0.46 | 1.47 | | | | |
| | | | C4 | 1.06 | 0.58 | 1.95 | | | | | 1.90 | 0.91 | 3.97 | | | | | 1.32 | 0.69 | 2.50 | | | | |
| M1 | | | | | | | | <0.0001 | 0.70 | 584 | | | | | <0.0001 | 0.76 | 563 | | | | | <0.0001 | 0.68 | 584 |
| | stage | | | | | | <0.0001 | | | | | | | <0.0001 | | | | | | | <0.0001 | | | |
| | | 1 | 2 | 2.55 | 1.20 | 5.40 | | | | | 2.79 | 0.63 | 12.41 | | | | | 1.74 | 0.77 | 3.95 | | | | |
| | | | 3 | 3.56 | 1.68 | 7.56 | | | | | 6.59 | 1.55 | 28.13 | | | | | 3.09 | 1.38 | 6.92 | | | | |
| | | | 4 | 12.99 | 6.15 | 27.43 | | | | | 26.44 | 6.35 | 110.05 | | | | | 7.53 | 3.35 | 16.92 | | | | |
| | microbial subtype | | | | | | 0.22 | | | | | | | 0.02 | | | | | | | 0.69 | | | |
| | | C1 | C2 | 0.90 | 0.63 | 1.29 | | | | | 1.47 | 0.88 | 2.46 | | | | | 1.12 | 0.73 | 1.71 | | | | |
| | | | C3 | 0.53 | 0.28 | 1.00 | | | | | 0.35 | 0.11 | 1.17 | | | | | 0.77 | 0.42 | 1.42 | | | | |
| | | | C4 | 0.93 | 0.49 | 1.75 | | | | | 1.77 | 0.81 | 3.87 | | | | | 1.16 | 0.58 | 2.35 | | | | |
| M2 | | | | | | | | <0.0001 | 0.71 | 567 | | | | | <0.0001 | 0.79 | 548 | | | | | <0.0001 | 0.74 | 567 |
| | stage | | | | | | <0.0001 | | | | | | | <0.0001 | | | | | | | <0.0001 | | | |
| | | 1 | 2 | 2.38 | 1.12 | 5.06 | | | | | 2.60 | 0.58 | 11.56 | | | | | 1.87 | 0.78 | 4.49 | | | | |
| | | | 3 | 3.72 | 1.75 | 7.92 | | | | | 7.47 | 1.75 | 31.94 | | | | | 4.29 | 1.81 | 10.20 | | | | |
| | | | 4 | 13.31 | 6.25 | 28.35 | | | | | 32.34 | 7.68 | 136.11 | | | | | 11.96 | 4.96 | 28.87 | | | | |
| | tumour site | | | | | | 0.12 | | | | | | | 0.003 | | | | | | | 0.01 | | | |
| | | proximal | distal | 0.72 | 0.49 | 1.07 | | | | | 0.44 | 0.25 | 0.79 | | | | | 0.54 | 0.33 | 0.86 | | | | |
| | | | rectal | 0.67 | 0.44 | 1.03 | | | | | 0.41 | 0.22 | 0.79 | | | | | 0.56 | 0.33 | 0.93 | | | | |
| | sex | | | | | | 0.44 | | | | | | | 0.86 | | | | | | | 0.80 | | | |
| | | male | female | 0.88 | 0.63 | 1.22 | | | | | 0.96 | 0.59 | 1.55 | | | | | 1.05 | 0.72 | 1.55 | | | | |
| | age | | | 1.00 | 0.99 | 1.02 | 0.64 | | | | 1.02 | 1.00 | 1.04 | 0.12 | | | | 1.04 | 1.02 | 1.05 | <0.0001 | | | |
| | microbial subtype | | | | | | 0.17 | | | | | | | 0.02 | | | | | | | 0.34 | | | |
| | | C1 | C2 | 0.86 | 0.59 | 1.25 | | | | | 1.28 | 0.74 | 2.19 | | | | | 0.93 | 0.60 | 1.46 | | | | |
| | | | C3 | 0.50 | 0.26 | 0.97 | | | | | 0.28 | 0.08 | 0.93 | | | | | 0.57 | 0.29 | 1.10 | | | | |
| | | | C4 | 0.87 | 0.46 | 1.65 | | | | | 1.47 | 0.67 | 3.26 | | | | | 1.05 | 0.51 | 2.15 | | | | |

**Supplementary Figure 5.** ***Univariate and multivariate Cox regression models for CRC patients of the TCGA-COAD-READ cohort.***

Univariate models were fitted for baseline clinical, demographic and pathological characteristics and microbial subtyping. Two multivariate models (M1 and M2) were fitted to adjust by baseline patient characteristics selected *a priori*. HRs, 95% CIs, p-values computed by loglikelihood ratio tests, *c*-indices and number of included patients were reported. Multivariate model M1 controlled for stage while multivariate model M2 additionally adjusted for age, sex and tumour location.
