## Supplementary material for "Identification and characterization of four bacteriome- and mycobiome-derived subtypes in tumour and adjacent mucosa tissue of colorectal cancer patients": Captions for supplementary tables

### Captions for Supplementary Tables for

**Supplementary Tables**

**Supplementary Table 1.** ***Patient samples included in the study.*** Samples included in the study were tallied by source/cohort and type of specimen. Samples labelled as "tumour" indicate primary tumour specimens as those collected from recurrence, metastasis or peripheral blood were excluded from the study.

**Supplementary Table 2.** ***Clinical, demographic and pathological characteristics of the cancer patients for the in-house ICL and TCGA-COAD-READ colorectal cohorts.*** Statistical significant differences by cohort were assessed using the $\chi 2$ and the Kruskal tests for categorical and continuous variables, respectively. P-values were not adjusted for multiple comparisons.

**Supplementary Table 3.** ***Clinical, demographic and pathological characteristics of the cancer patients for the TCGA pan-cancer cohorts.*** Statistical significant differences by cohort were assessed using the $\chi 2$ and the Kruskal tests for categorical and continuous variables, respectively. P-values were not adjusted for multiple comparisons. The TCGA-COAD-READ cohort (colorectal cancer cohort) was included for completeness.

**Supplementary Table 4.** ***Association between microbial subtyping and bacterial and fungal families.*** Statistically significant differences in relative abundance of bacteria and fungi at the family taxonomic rank by microbial subtype were assessed by ANOVA tests followed by pairwise post-hoc Tukey tests.

**Supplementary Table 5.** ***Clinical and demographic characteristics of the healthy patients with colon-sigmoid or colon-transverse samples from the GTEx cohorts.*** Statistical significant differences by cohort were assessed using the $\chi 2$ and the Kruskal tests for categorical and continuous variables, respectively. P-values were not adjusted for multiple comparisons.

**Supplementary Table 6.** ***Association between mutational status and microbial subtypes in the TCGA-COAD-READ patients.*** Statistical significance was assessed by $\chi 2$ independence tests and unadjusted- and Benjamini-Hochberg FDR-corrected mod-likelihood P-values are reported for each mutation that was found to be statistically significantly altered.

**Supplementary Table 7.** ***Association between gene expression profiles and microbial subtyping in the TCGA-COAD-READ patients.*** Statistical significance was assessed by Kruskal-Wallis H-tests and unadjusted- and Benjamini-Hochberg FDR-corrected P-values are reported for each of the tested genes. Analysis was restricted to the top 5000 most variant genes.

**Supplementary Table 8.** ***Association between protein expression profiles determined by Reverse Phase Protein Array (RPPA) and microbial subtyping in the TCGA-COAD-READ patients.*** Statistical significance was assessed by Kruskal-Wallis H-tests and unadjusted- and Benjamini-Hochberg FDR-corrected P-values are reported for each of the tested genes. Analysis was restricted to the top 5000 most variant genes.

**Supplementary Table 9.** ***Clinical, demographic and pathological characteristics of the cancer patients for the TCGA-COAD-READ colorectal cohort stratified by microbial subtype.*** Statistical significant differences by microbial subtyping were assessed using the $\chi 2$ and the Kruskal tests for categorical and continuous variables, respectively. P-values were not adjusted for multiple comparisons.

**Supplementary Table 10.** ***Association between gene expression profiles and microbial subtyping in the TCGA pan-cancer patient cohorts.*** Statistical significance was assessed by Kruskal-Wallis H-tests and unadjusted- and FDR-corrected P-values are reported for each of the tested genes. Analysis was restricted to the top 5000 most variant genes.

**Supplementary Table 11.** ***Association between protein expression profiles determined by Reverse Phase Protein Array (RPPA) and microbial subtyping in the TCGA pan-cancer patient cohorts.*** Statistical significance was assessed by Kruskal-Wallis H-tests and unadjusted- and FDR-corrected P-values are reported for each of the tested genes. Analysis was restricted to the top 5000 most variant genes.

**Supplementary Table 12.** ***Univariate Cox regression models by microbial subtyping stratified by cancer type.*** Univariate Cox regression models by microbial subtyping for DFS, DSS and OS endpoints for each individual cancer type. Analysis was restricted to cancer types where at least 2 microbial subgroups were identified with at least 10 patients each.
